# Recognition of N6-methyladenosine by the YTHDC1 YTH domain studied by molecular dynamics and NMR spectroscopy: The role of hydration

**DOI:** 10.1101/2021.02.08.430239

**Authors:** Miroslav Krepl, Fred Franz Damberger, Christine von Schroetter, Dominik Theler, Pavlína Pokorná, Frédéric H.-T. Allain, Jiří Šponer

**Affiliations:** Institute of Biophysics of the Czech Academy of Sciences, Kralovopolska 135, 612 65 Brno, Czech Republic; Department of Biology, ETH Zürich, Institute of Biochemistry, 8093 Zürich, Switzerland; National Centre for Biomolecular Research, Faculty of Science, Masaryk University, Kamenice 5, 625 00 Brno, Czech Republic; Regional Centre of Advanced Technologies and Materials, Czech Advanced Technology and Research Institute (CATRIN), Palacký University Olomouc, Olomouc 783 71, Czech Republic

**Keywords:** YTH domain, YTHDC1, water-bridges, hydration, molecular dynamics, N6-methyladenosine, NMR spectroscopy

## Abstract

The YTH domain of YTHDC1 belongs to a class of protein “readers”, recognizing the N6-methyladenosine (m^6^A) chemical modification in mRNA. Static ensemble-averaged structures revealed details of N6-methyl recognition via a conserved aromatic cage. Here, we performed molecular dynamics (MD) simulations along with nuclear magnetic resonance (NMR) and isothermal titration calorimetry (ITC) to examine how dynamics and solvent interactions contribute to the m^6^A recognition and negative selectivity towards unmethylated substrate. The structured water molecules surrounding the bound RNA and the methylated substrate’s ability to exclude bulk water molecules contribute to the YTH domain’s preference for m^6^A. Intrusions of bulk water deep into the binding pocket disrupt binding of unmethylated adenosine. The YTHDC1’s preference for the 5′-Gm^6^A-3′ motif is partially facilitated by a network of water-mediated interactions between the 2-amino group of the guanosine and residues in the m^6^A binding pocket. The 5′-Im^6^A-3′ (where I is inosine) motif can be recognized too but disruption of the water network lowers affinity. The D479A mutant also disrupts the water network and destabilizes m^6^A binding. Our interdisciplinary study of YTHDC1 protein/RNA complex reveals an unusual physical mechanism by which solvent interactions contributes towards m^6^A recognition.

## Introduction

N6-methyladenosine (m^6^A) is the most common posttranscriptional modification of nucleobases in mRNA.^1-2^ It can influence basic cellular processes such as cell differentiation and proliferation by changing the stability of specific RNA transcripts or by promoting their translation.^3-4^ It is most often localized near stop-codons or in 5′- or 3′-untranslated regions of mRNA.^5^ To handle the full cycle of the m^6^A metabolism, the cell requires proteins that can introduce, detect, and remove this modification.^6-8^ These proteins are commonly known as writers, readers and erasers, respectively. Among the readers, the YT521-B homology (YTH) family of proteins is the most prominent. The YTH domain is evolutionary conserved in eukaryotes, possessing a characteristic α_1_β_1_α_2_β_2_α_3_β_3_β_4_β_5_β_6_α_4_ secondary structure.^3, 9^ It facilitates recognition of the methyl group in N6-methyladenosine by a set of aromatic amino acids, including two tryptophans which are absolutely conserved in all YTH domains.^10^ These residues form an aromatic cage which binds N6-methyladenosine with 50-fold higher affinity than adenosine (Figure 1). It was suggested that this preference is based on a set of van der Waals (vdW) interactions between the residues lining the m^6^A binding pocket (including the conserved tryptophans) and the N6-methyl group.^10^ Structurally, the aromatic cage is similar to the binding sites of methylated lysine observed in the evolutionary unrelated chromo and tudor protein domains,^11^ making it a good example of convergent evolution in recognition of methylated modifications of biomolecules, be they proteins or nucleic acids. Here, we explore the YTH domain of YTHDC1 protein which facilitates the regulation of pre-mRNA splicing by recruiting splicing factors.^12^ The YTHDC1 protein interacts with RNA solely by its single YTH domain.^9^ Unusually, the YTHDC1 preferentially recognizes 5’-Gm^6^A-3’ motifs in RNA.^13-14^ The other YTH-domain-containing proteins, such as the YTHDF family,^15^ do not display any preference for nucleotide preceding m^6^A.

**Figure 1.**
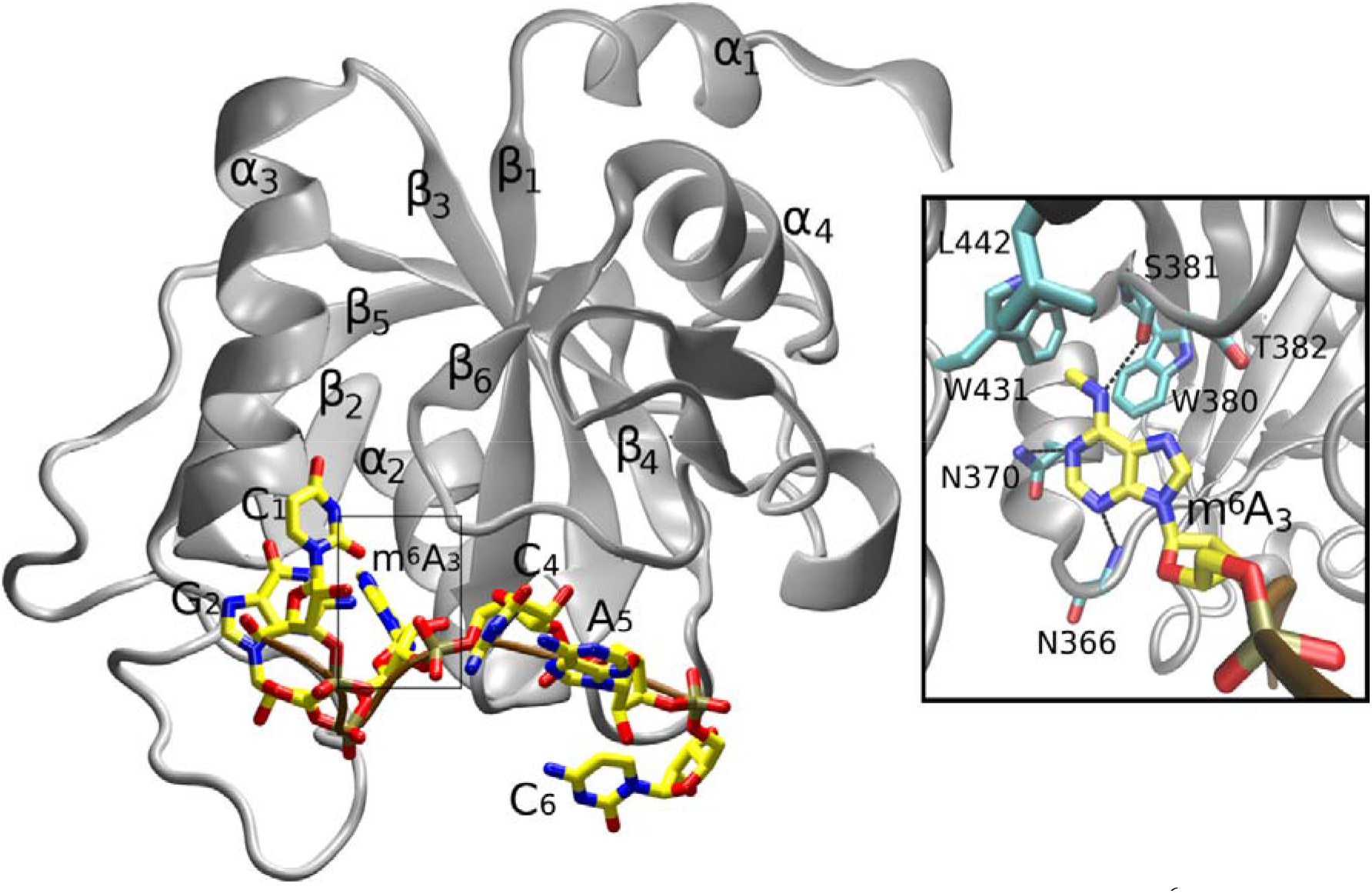
YTH domain of the YTHDC1 protein complexed with a 5′-C_1_G_2_m^6^A_3_C_4_A_5_C_6_-3′ RNA oligonucleotide.^13^ Detail of the m^6^A binding pocket is shown in the inset. Secondary structure of the protein is labeled and shown as gray ribbon. The RNA backbone is in brown. The carbon atoms are colored yellow and cyan in RNA and protein, respectively, whereas nitrogen, oxygen and phosphorus are blue, red, and gold. Protein/RNA H-bonds are marked with dashed black lines. Unless specified otherwise, same color coding of atoms and marking for H-bonds were used in the subsequent figures.

We use molecular dynamics (MD) simulations to study the YTH domain of the YTHDC1 protein, both in isolation and bound to RNA oligonucleotide containing the 5’-Gm^6^A-3’ motif (Figure 1). MD describes the Boltzmann distribution (populations) of molecular conformations using carefully calibrated empirical models – molecular mechanical force fields. In the past, the MD simulations have been successfully applied many times in studies of protein/RNA complexes,^16^ including the YTH domain.^14, 17-18^ By providing essentially infinite temporal and spatial resolution, the MD can extend upon the information from ensemble-averaged experimental data. It is particularly well-suited for studies of hydration and its structural role in biomolecular complexes.^19-20^ The main limitations of MD are the quality of the force field and the affordable timescale (sampling) of the simulation,^16^ necessitating careful interpretation of the simulation data. Thus we apply NMR spectroscopy experiments and isothermal titration calorimetry (ITC) measurements to verify and corroborate the most important results predicted by our simulations.^21^

By elucidating the role of dynamics and solvent interactions, our data provides a more complete picture of the YTH domain preference for m^6^A over A, going beyond the static structural details revealed by experimental data.^13, 22-23^ In addition to the absence of stabilizing vdW interactions with the protein, the unmethylated adenosine is more attractive for water and is also sterically unable to fully displace bulk water molecules from the m^6^A binding pocket, which increases the frustration of interactions at the protein/RNA interface. This disrupts its binding and could facilitate fast turnover of incorrect substrates. The preference for the 5’-Gm^6^A-3’ motif, which is unique to the YTHDC1 among the other YTH domains, is primarily facilitated by direct protein/RNA H-bonds with the guanosine (henceforth referred to as G_2_), as shown by the experimental structures.^13, 24^ Here, our calculations reveal the G_2_ is also involved in two guanine-specific water-bridge interactions via its 2-amino group. These water-bridge interactions connect the G_2_ to a water network within the m^6^A binding pocket. Mutating the G_2_ into I_2_ (inosine) abolishes this connection, impacting residues deep in the m^6^A binding pocket and slightly reducing the protein/RNA binding affinity. We quantify the overall change in free energy of RNA binding by thermodynamic integration (TI) free-energy calculations and ITC experimental measurements and find good agreement. The water network can be even more disrupted by a D479A mutation in the protein.

## Material and Methods

### System building

We used structures of the free YTHDC1 (PDB ID: 4R3H)^22^ and its complex with 5′-CGm^6^ACAC-3′ RNA (PDB ID: 2MTV)^13^ as starting structures for all our MD simulations. Only the first conformer of the 2MTV ensemble was used as all the conformers are very similar due to relative abundance of the NMR signals. The initial files were prepared by the xLeap module of AMBER 18.^25^ All experimentally determined atoms were utilized. The missing residue side-chains in the structure of the free YTHDC1 were added by xLeap and the structure of the missing β4/β5 protein loop was taken from another structure of the YTHDC1 complexed with RNA (PDB ID: 4R3I).^22^ Both the 4r3i and 2mtv structures contain 5’-Gm^6^A-3’ motif in the bound oligonucleotide and show a virtually identical protein/RNA interface. For our simulations of the protein/RNA complex, we have utilized the 2mtv structure which was more convenient for this study but we also utilize the X-ray structure^22^ for interpreting our results (see Supporting Information for details). We used the ff12SB^26^ and bsc0χ_OL3_ (i.e., OL3)^27-28^ force fields to describe the protein and RNA, respectively; see Refs.^16, 29^ for reasons for choosing the ff12SB version of the protein force field. The electrostatic potentials of the N6,N1-dimethylated adenine base and of the methylamine were computed by Gaussian 09^30^ using HF/6-31G* level of theory. The partial charges of the m^6^A base and of the methylamine were subsequently derived via the RESP method.^31^ The remaining force-field parameters of m^6^A were taken from the standard OL3 RNA force field and the parameters of the methylamine were adapted from parameters of the N6-methyl segment of the base. The partial charges and parameters of inosine were taken from Ref.^32^ and OL3 parameters for guanosine^28^ were used to describe its χ dihedral. In all simulations, the biomolecules were surrounded in an octahedral box of SPC/E^33^ water molecules with minimal distance of 12 Å between the solute and the box border. Some simulations were also done with OPC waters and modified vdW parameters of phosphates supplemented by modified RNA backbone dihedrals (abbreviated as χ_OL3__CP(OPC) force-field combination).^34-36^ KCl ions^37^ were added to neutralize all systems and to achieve an excess salt concentration of 0.15 M. These solvent conditions work well for MD simulations of protein-RNA complexes.^16^ Note that, inevitably, the buffer used in the NMR experiments is different from the simple salt conditions in MD simulations due to inherent differences between the two methods. It should, however, not preclude a comparison of the experimental data with the simulations. For extensive discussion of differences between the buffers used in NMR studies of protein-RNA complexes and simulation buffers, please see Supplementary of Ref.^38^.

### Simulation protocol

The initial minimization and equilibration of the systems was performed in the pmemd.MPI implementation of AMBER according to the protocol described in Ref.^39^. In simulations of the protein/RNA complex structure, this was followed by 120 ns of simulations where we gradually released the experimental NMR-restraints. This protocol was designed to prevent abrupt departures from experimental geometry which is known to sometimes occur in early stages of simulations of NMR structures, as described in detail elsewhere.^38^ Afterwards, the production simulations were run using the GPU-based pmemd.cuda^40^ on GTX 1080 graphic card, obtaining trajectories at a speed of ca 260 ns per day. In the preliminary set of simulations, we observed reversible perturbations of the native protein/RNA interactions for the C_4_, A_5_, and C_6_ which, in some cases, led to loss of the protein/RNA interface for these peripheral nucleotides, but not the G_2_ and m^6^A_3_ which were of our primary interest. Nevertheless, to avoid the potential sampling uncertainties when interpreting the data for the G_2_ and m^6^A_3_ in such trajectories, we used mild 1 kcal/mol HBfix potential^41^ to increase stability of the native A_5_(OP1)/K364(NZ) and C_6_(OP1)/K475(NZ) interactions in all subsequent simulations; see Ref.^42^ for justifications of such approach in simulations of protein/RNA complexes. HBfix was originally suggested as a simple short-distance force-field term that introduces a gentle stabilizing (or destabilizing) potential between two atoms with non-zero forces limited to a distance range relevant for H-bond interaction.^41^ It can, for example, mimic an induction term.

In some simulations, we used NOEfix (first tested in this work) instead of HBfix to increase the simulation agreement with experiment. In the NOEfix, the HBfix potential function is modified to accommodate the NOE upper-bound distances instead of H-bonds for few carefully selected experimental NOEs measured by NMR. In contrast to standard restraints, NOEfix does not introduce a continuously growing penalty with increasing distances. In other words, it allows smooth sampling of substates that temporarily or even permanently violate NOEs. Although not a panacea, NOEfix potential offers an alternative and easily applicable tool for curation of MD simulation trajectories based on the NMR data and known deficiencies of the force fields. Both structure-specific HBfix and NOEfix produced highly similar results for simulations of the YTHDC1 system, cross-validating the two approaches; see Supporting Information for further details. In all simulations, we used the SHAKE algorithm^43^ and hydrogen mass repartitioning,^44^ allowing a 4 fs integration step. Particle mesh Ewald summation^45^ was used to describe electrostatic interactions and periodic boundary conditions were applied to handle system border bias. The cut-off distance for Lennard-Jones interactions was 9 Å. A Langevin thermostat and a Monte Carlo barostat^25^ were used to regulate temperature and pressure, respectively.

### Thermodynamic integration

We used a mixed single/double topology approach as implemented in AMBER 18 (including update 16) to set up the Thermodynamic integration (TI) calculations. Specifically, a segment of the mutated residue undergoing the alchemical transformation was represented by a dual topology model while the rest of the molecule was described by single topology. We used the soft-core vdW potentials^46^ to describe the appearing and disappearing atoms in the dual topology region. Fully unrestrained simulations were carried out for nine lambda windows in parallel runs. The length of the individual lambda windows was 20 and 100 ns for reference RNA single-strand and protein/RNA complex simulations, respectively. The length of the complex simulations was greater to increase sampling of the conformational and water-network changes at the protein/RNA interface induced by the alchemical transformation. See Supplementary for further details and for the TI calculations of solvation energies of free m^6^A and A nucleobases.

### Protein and RNA Preparation

The ^15^N-labeled YTH domain of *R. Norvegicus* YTHDC1 (residues 347-502) was prepared as previously described^13^ except that for the size exclusion chromatography (SEC) a buffer consisting of 25 mM NaH_2_PO_4_, 25 mM NaCl and 10 mM β-Mercaptoethanol at pH 7 (pH adjusted with NaOH) was used and 880 units per ml SEC sample RNAseOUT(Invitrogen) as RNAse inhibitor were added to the sample prior to the size exclusion chromatography purification step. Note that in Ref.^13^, it was erroneously reported that the SEC buffer was prepared using NaH_2_PO_4_ instead of Na_2_HPO_4._ RNAs were obtained from Horizon Discovery Ltd, deprotected according to the supplier’s instructions, lyophilized and resuspended in the SEC buffer.

### Isothermal titration calorimetry (ITC)

Protein and RNA were dialyzed in SEC buffer before ITC measurements. ITC measurements were conducted at 20°C on a VP-ITC calorimeter (MicroCal) with a protein concentration of 10 μM in the cell and with syringe concentrations of 117 uM UGm^6^ACAC and 124 uM UIm^6^ACAC, respectively. An initial injection of 2 μl was followed by 45 injections of 6 μl each with an interval of 300 s between injections and a stirring speed of 307 rpm. Data analysis was perfomed with the Origin 7.0 software.

### NMR

NMR measurements were conducted in the SEC buffer supplemented with 10% D_2_O as lock substance. NMR spectra were obtained on an AVIII 600MHz spectrometer equipped with a cryoprobe at 293K. Spectra were processed using Topspin3 (Bruker) and analysed with CARA^47^ (www.nmr.ch). Titrations with RNA were performed by adding concentrated aliquots in NMR buffer to 0.9 mM YTHDC1 samples which showed slow exchange behavior with separate signals observed for the free and RNA-bound states for both I_2_m^6^A_3_ and G_2_m^6^A_3_ RNAs. Assignments for the amides of YTHDC1 in the I_2_m^6^A_3_ complex were obtained from a 3D ^15^N-resolved NOESY^48^ (in-house implementation) of the complex by adjusting the position of H,N correlations so that NOESY towers matched the patterns observed in the G_2_m^6^A_3_ complex. This was possible thanks to the highly similar aliphatic ^1^H chemical shifts and NOESY towers in both structures (Supplementary Figure S1). Final H,N peak positions used for obtaining chemical shift perturbations (CSPs) were then adjusted to match ^15^N,^1^H-HSQCs of the 1:1 G_2_m^6^A_3_ and I_2_m^6^A_3_ complexes. CSPs were calculated with the formula CSP=[(0.2Δδ_N_)^2^+ Δδ_H_^2^]^1/2^. Where Δδ_N_ and Δδ_H_ are the chemical shift differences of the two complexes for backbone amide ^15^N and ^1^H resonances, respectively.

### Analyses

We have used cpptraj and VMD for ensemble analysis and visual inspection of the MD simulation trajectories, respectively^49-50^ Raster3D, Povray and Gnuplot were used to prepare figures and graphs.^51^ H-bonds were analyzed by monitoring atom distances between heavy atom donors and acceptors and donor-hydrogen-acceptor angles. The force-field performance of the protein/RNA complex simulations was evaluated by computing the 1/r^6^ averaged distance violations of the NOEs in the simulation ensemble by sander and comparing them with experimentally measured upper bounds.^38^ The NOE distance in MD simulations was considered satisfied when its average value did not exceed the experimental upper bound distance plus 0.3 u. The ion binding sites in simulations were evaluated by computing the K^+^ (or Cl^-^) ion density grid and by analysis of the interactions between ions and individual solute atoms.

## Results

### Overview of the results

We report a series of MD simulations (Table 1) of the YTH domain of YTHDC1 bound to m-^6^A or standard adenosine RNA oligonucleotides to examine how dynamics in the complex is affected by the N6-methyl modification. We also simulated the free protein to characterize structure and dynamics of the m^6^A binding pocket when no nucleotide is bound. The simulations with m^6^A_3_ revealed a network of structured water molecules and we subsequently performed simulations with an I_2_ nucleotide present instead of G_2_ to show how loss of the water interactions coordinated by the guanine’s amino group affects the protein/RNA interface. The effects of binding I_2_ or G_2_-containing RNAs to YTH are also examined by NMR and ITC measurements. Lastly, we examine the dynamics of the m^6^A_3_ RNA substrate complexed with YTH D479A mutant which severely disrupts the water interaction network.

**Table 1.** List of simulations.^a^

| simulated system | number of simulations × length [μs] |
| --- | --- |
| 2mtv_m<sup>6</sup>A<sub>3</sub> | 8 × 1 μs |
| 2mtv_m<sup>6</sup>A<sub>3</sub>_NOEfix312<sup>b</sup> | 6 × 1 μs |
| 2mtv_m<sup>6</sup>A<sub>3</sub>_NOEfix18<sup>b</sup> | 6 × 1 μs |
| 2mtv_m<sup>6</sup>A<sub>3</sub>_CP<sup>c</sup> | 5 × 1 μs |
| 2mtv_A<sub>3</sub> | 4 × 1 μs |
| 2mtv_I<sub>2</sub> | 4 × 1 μs |
| 2mtv_D479A_m<sup>6</sup>A<sub>3</sub> | 4 × 1 μs |
| 4r3h | 4 × 1 μs |
| 4r3h_methylamine | 2 × 1 μs |
| 2mtv_TI_m<sup>6</sup>A<sub>3</sub>/A<sub>3</sub> | 54 × 100 ns, 54 × 20 ns |
| 2mtv_TI_G<sub>2</sub>/I<sub>2</sub> | 54 × 100 ns, 54 × 20 ns |

<sup>a</sup>Experimental structures of the free protein (PDB ID: 4R3H)<sup>22</sup> or the protein/RNA complex (PDB ID: 2MTV)<sup>13</sup> served as starting structures in simulations. The m<sup>6</sup>A<sub>3</sub> and A<sub>3</sub> indicate that the N6-methyladenosine and standard adenosine, respectively, were the third RNA nucleotide. I<sub>2</sub> indicates a G to I substitution of the second nucleotide. D479A indicates that the aspartate 479 of YTH was mutated into alanine. Methylamine indicates that twenty methylamines were added into the simulation water box before start. <sup>b</sup>NOEfix function was applied to all (312) or selected (18) experimental NOE distances showing ensemble-averaged distance violations in the 2mtv\_m<sup>6</sup>A<sub>3</sub> simulations. <sup>c</sup>Modified vdW parameters of the phosphates<sup>35</sup> with associated modified RNA backbone dihedral parameters<sup>36</sup> and OPC water model<sup>34</sup> were used.

### The very stable interaction of N6-methylated adenosine with YTHDC1 is mediated by H-bonding, stacking and water bridges

Experimental structures of the YTHDC1 complex^13, 22^ show m^6^A_3_ base forming stacking and H-bonding interactions with the protein. Namely, there are m^6^A_3_(N6)/S381(O), m^6^A_3_(N1)/N370(ND2), and m^6^A_3_(N3)/N366(N) H-bonds. In addition, one face of the base and the N6-methyl group stack with W380 and W431 side-chains, respectively, and the other face of the base stacks with the side-chain of L442 (Figure 1). These interactions were entirely stable in all simulations. The simulations revealed a network of structured water molecules (water bridges) facilitating additional protein/RNA interactions. Namely, there was a highly occupied water bridge involving four solute atoms – m^6^A_3_(N7), W380(NE1), T382(OG1), and D479(OD) (Figure 2), having water exchange time with bulk on the scale of hundreds of nanoseconds (Table 2). Location of this hydration site is in excellent agreement with the corresponding X-ray structure (PDB ID: 4R3I).^22^ The simulations also revealed a water bridge between T382(OG1) and D479(OD) and another between m^6^A_3_(O2’) and K364(O) (Figure 2). These water molecules are not visible in the X-ray structure, likely due to the conformational flexibility of this region, and were thus uniquely predicted by the MD simulations with an exchange timescale of tens of nanoseconds. Overall, the simulations predict an intricate network of water molecules within and near the entrance to the m^6^A_3_ binding pocket (Figure 2); for discussion of diverse long-residency hydration sites seen in MD simulations in RNA and protein-RNA systems see Ref.^16^. The NMR experimental structure we used as starting structure in our protein/RNA simulations does not provide any *a priori* information about the distribution of solvent molecules and all observed hydration sites were thus spontaneously established during the equilibration or early stages of the simulations. Notably, we observed the same water network near the m^6^A_3_ binding pocket also while using the OPC water model (see Methods) so the results do not depend on the utilized water model. Lastly, a potassium binding site was transiently observed in all MD simulations, coordinated by amino acids from the β2, α3, α5 segments (Supplementary Figure S2). The site had a relatively low average occupancy in simulations of only ca 25%, indicating that structural ions do not play a prominent role in stabilization of this complex.

**Table 2.** Occupancies and maximum observed lifetimes of the structured water molecules in MD simulations of the individual systems.^a^

| water molecule / simulation ensemble | occupancies (in %) |  |  |  |  |  |
| --- | --- | --- | --- | --- | --- | --- |
|  | w1 | w2 | w3 | w4 | w5 | w6 |
| 2mtv_$m^6A_3$ | 90 | 52 | 53 | 42 | 50 | - |
| 2mtv_$m^6A_3$_NOEfix312 | 90 | 71 | 55 | 55 | 68 | - |
| 2mtv_$m^6A_3$_NOEfix18 | 86 | 55 | 57 | 52 | 45 | - |
| **2mtv_A<sub>3</sub><sup>b</sup>** | 84 | 46 | 59 | 41 | 52 | - |
| **2mtv_I<sub>2</sub>** | 97 | 80 | - | - | - | 69 |
| **2mtv_D479A_m<sup>6</sup>A<sub>3</sub>** | 60 | - | - | 59 | 49 | - |
| **lifetimes (in ns)** |  |  |  |  |  |  |
|  | **w1** | **w2** | **w3** | **w4** | **w5** | **w6** |
| **2mtv_m<sup>6</sup>A<sub>3</sub>** | 660 ± 166 | 446 ± 253 | 280 ± 146 | 37 ± 35 | 31 ± 46 | - |
| **2mtv_m<sup>6</sup>A<sub>3</sub>_NOEfix312** | 624 ± 129 | 426 ± 69 | 121 ± 130 | 29 ± 32 | 44 ± 50 | - |
| **2mtv_m<sup>6</sup>A<sub>3</sub>_NOEfix18** | 604 ± 162 | 411 ± 122 | 156 ± 161 | 26 ± 38 | 55 ± 37 | - |
| **2mtv_A<sub>3</sub><sup>b</sup>** | 363 ± 324 | 148 ± 65 | 186 ± 97 | 33 ± 32 | 23 ± 23 | - |
| **2mtv_I<sub>2</sub>** | 703 ± 66 | 444 ± 35 | - | - | - | 80 ± 37 |
| **2mtv_D479A_m<sup>6</sup>A<sub>3</sub>** | 418 ± 250 | - | - | 37 ± 21 | 7 ± 8 | - |

<sup>a</sup>Designations of the individual water molecules are the same as in Figure 6. The cells marked with “-” indicate that the water bridge was not formed in the specified simulation ensemble. Occupancies indicate percentages of the simulation ensemble in which the specified water bridges were present. An individual water bridge was considered present when its water molecule participated in H-bonding interactions as shown in Figure 6. Lifetimes indicate maximum time a single water molecule was observed forming the individual water bridges as shown in Figure 6, averaged over the individual simulations of the ensemble. Individual simulations were taken as samples for calculating the standard deviation. Note that in some trajectories, the w1 and w2 water did not exchange on the simulation timescale after being first established. <sup>b</sup>Simulation frames where the A<sub>3</sub> nucleotide temporarily or permanently drifted out of its binding pocket (see above and Figure 3) were not included in the analyses.

**Figure 2.**
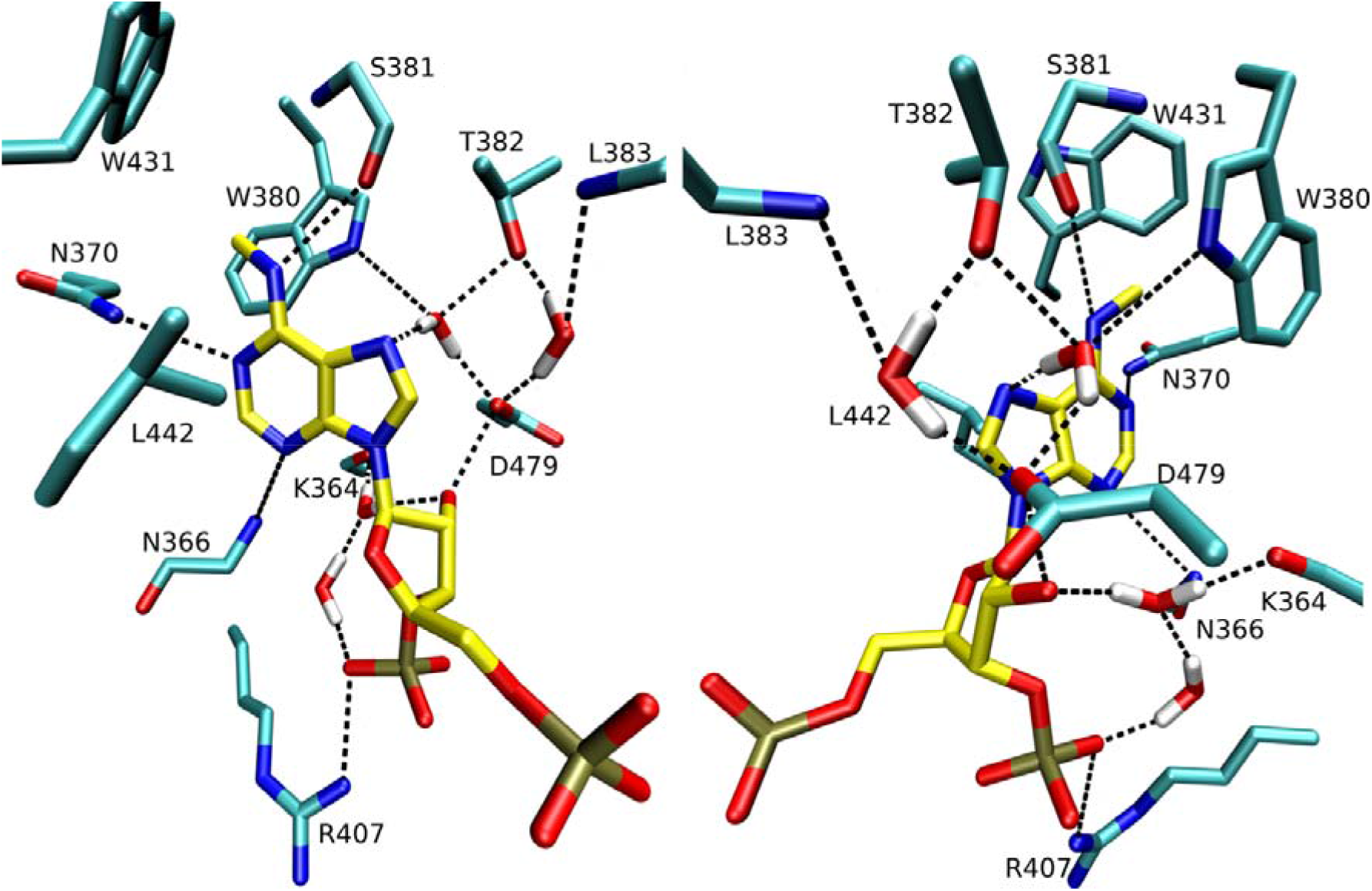
Recognition of the m^6^A_3_ nucleotide in MD simulations shown from two different angles. The participating amino-acids are labeled. Solvent hydrogens are white. For clarity, the solute hydrogens are not displayed.

### Unmodified adenosine permits invasions of water molecules into the m^6^A_3_ binding pocket

There are no experimental structures of YTH domain bound to standard adenosine, even though it is known to bind in the same manner as m^6^A albeit with lower affinity.^13^ Here, we have performed simulations of the protein/RNA complex with A_3_, with starting structure obtained by substituting the N6-methyl group with hydrogen. These simulations at first showed an identical development as for the m^6^A_3_ variant, with all H-bonds and stacking interactions preserved, except those formed by the N6-methyl group. The same water bridges near the binding pocket were formed with slightly altered lifetimes (Table 2). However, all simulations subsequently showed periodic intrusions of a single water molecule from outside the pocket which could temporarily enter the region between the adenosine and the side-chain of W431 (Figure 3A). On average, a water molecule was present at this location in ∼9% of the simulation frames. The same location is occupied by N6-methyl group in the m^6^A_3_ systems (Figure 2). We interpret these water intrusions to be a direct consequence of the methyl group’s removal, resulting in a void which cannot be easily eliminated by reorientation of atoms forming the binding pocket. During its intrusions, the single water molecule formed one H-bond with the H61 atom of A_3_ (Figure 3A) but since there were no other hydrogen bond donors or acceptors nearby, it was unstable in this position and was soon expelled again from the pocket. Eventually, another water molecule would come in, repeating the process. On average, in our simulation set we observed ∼90 water molecule exchanges per 1 µs in this position.

**Figure 3.**
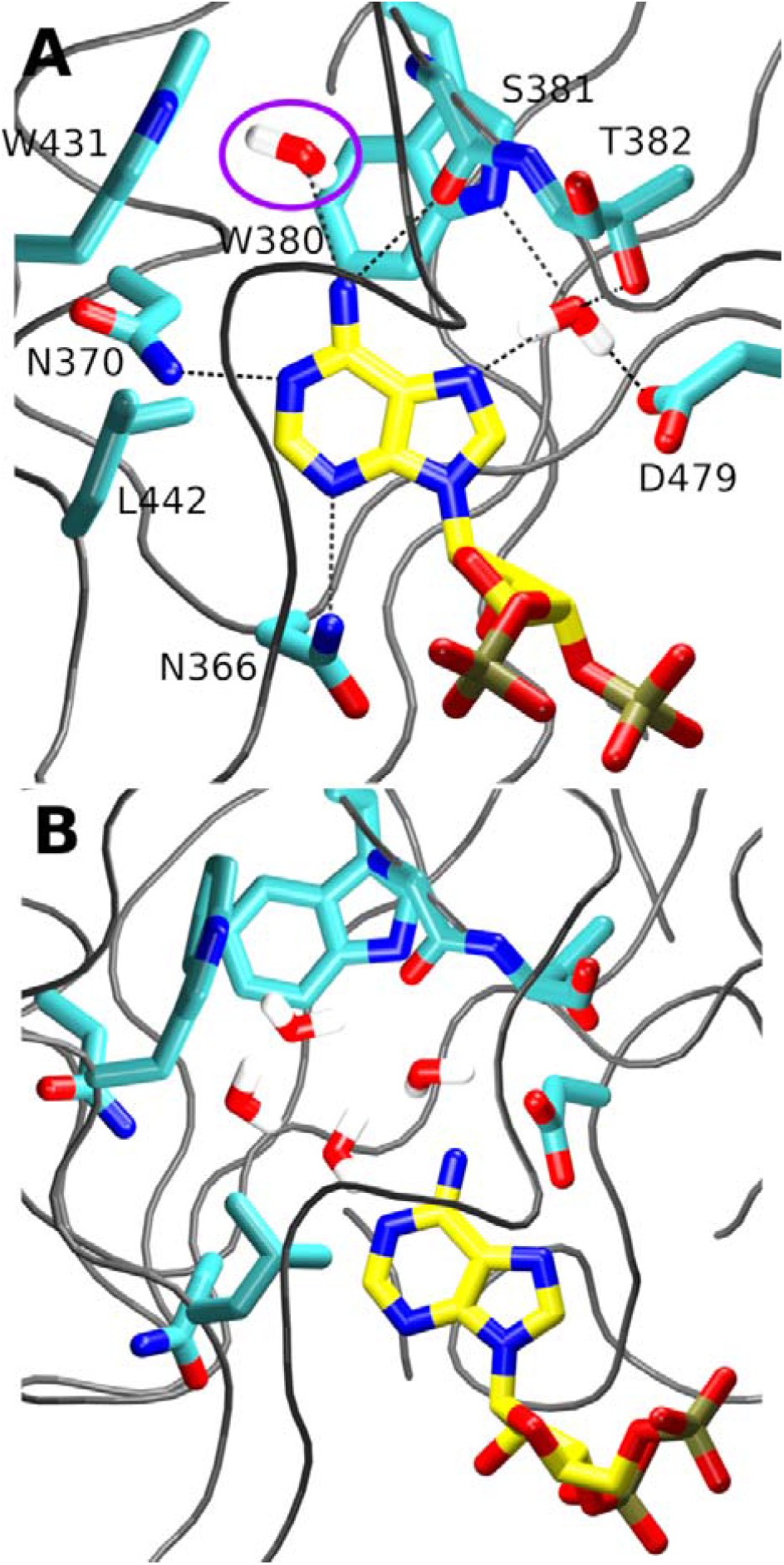
(**A**) A single water molecule (violet circle) periodically invaded the binding pocket of the A_3_ nucleotide in MD simulations of the 2mtv_A_3_ system, occupying the space where the N6-methyl group of the m^6^A_3_ nucleotide would normally be located. (**B**) In two simulations, the intrusion of the first water molecule attracted additional waters, in one case leading to complete expulsion of the A_3_ nucleotide from its binding pocket.

In two simulations, we also observed that instead of being expelled, the single water molecule gradually attracted additional water molecules from the bulk solvent into the binding pocket (Figure 3B). In the two simulations where this happened, the distance between A_3_ and the W431 side-chain started to increase as a result and the adenosine gradually drifted out of its binding pocket in order to make room for the incoming water molecules. In the first of the two simulations, this process was reversible on our simulation timescale. In the second, we observed a complete expulsion of A_3_ from its binding pocket facilitated by these invading water molecules after ca 800 ns of simulation.

### Thermodynamic integration reproduces lower binding affinity of standard adenosine

Earlier experiments have reported ∼50 fold stronger binding (free-energy difference of ∼2.3 kcal/mol) of m^6^A_3_ to the YTH domain compared to A_3_ as well as highly similar chemical shifts in presence of both ligands, suggesting they are bound identically.^13^ At first sight, this would make the system an ideal target for alchemical free energy calculations as the ligand transformation is likely to be sufficiently sampled even on relatively short simulation timescales. Indeed, a recent computational study of this system (using a different force field) utilized thermodynamic integration (TI) and indicated 1.7 kcal/mol penalty for binding of the unmethylated substrate.^18^ However, the calculations were performed for a model system with an isolated m^6^A nucleoside rather than a full oligonucleotide and the solute atoms were immobilized by positional restraints.^18^ Furthermore, our MD simulations indicated that a single water molecule often invades the binding pocket in presence of standard adenosine (see above and Figure 3) which somewhat complicates the sampling of m^6^A_3_→A_3_ TI calculations; see Supporting Information for further details.

To account for the invading water, we have performed two sets of TI calculations based on two different starting geometries – the forward one starting from a structure where the m^6^A_3_ nucleotide is bound in its binding pocket as observed in the experimental structure; and a backward calculation, starting from a simulation snapshot of an A_3_ system with a single water molecule already present in the binding pocket (Figure 3). Note that although not strictly necessary, we performed both forward and backward calculations also for the single-stranded RNAs. Finally, these forward and backward calculations yielded ΔΔG values of 4.3 ± 0.5 and −2.9 ± 0.4 kcal/mol, respectively. Thus, on average the calculations predicted a free-energy penalty of 3.6 ± 0.9 kcal/mol for binding of the A_3_ to the YTH domain compared to m^6^A_3_. The TI slightly overestimated the effects of the m^6^A_3_/A_3_ substitution. A possible reason for this is that the involvement of the solvent molecules was still insufficiently accounted for by our simulation setup. This is supported by the fact that the backward calculation was much closer to the experimentally observed value than the forward calculation. Nevertheless, considering the problems and inherent inaccuracies of free-energy calculation methods (stemming from both sampling limitations and molecular mechanics^16, 52^) we suggest that results of our TI calculations are still in relatively good agreement with the experiment.

### High solvation free energy of A_3_ compromises its ability to displace waters from within the binding pocket

The unmethylated adenosine was unable to prevent water molecules from entering the binding pocket (see above and Figure 3). We hypothesized that it could also be less effective in displacing the water molecules present inside the binding pocket during the RNA binding. The need to displace these water molecules was demonstrated by our simulations of the free YTHDC1 protein (Table 1)^22^ which showed that even though the side-chains forming the m^6^A_3_ binding pocket exhibited larger fluctuations than in the complex, the pocket was essentially pre-formed in the free protein. It was also filled with water molecules at all times, with rapid sub-ns exchange with the bulk solvent (Figure 4). This indicates that during RNA binding, the incoming nucleotide must displace these water molecules from within the pocket. Our TI calculations indicated that the free energy of solvation of the free A_3_ base is 0.67 kcal/mol more favorable than for the m^6^A_3_ (Supporting Information). Therefore, displacement of water molecules from within the binding pocket is less favorable process with A_3_ to begin with. It is further amplified by the fact that the protein is unable to form proper compensatory interactions with the H61 atom in place of the displaced water molecules (see above and Figure 3). We suggest the difference in the solvation energy of the two bases is thus straightforwardly contributing to their different binding affinity towards YTHDC1. The rest is made up by the favorable interactions between the protein and the N6-methyl group present solely with the m^6^A_3_.

**Figure 4.**
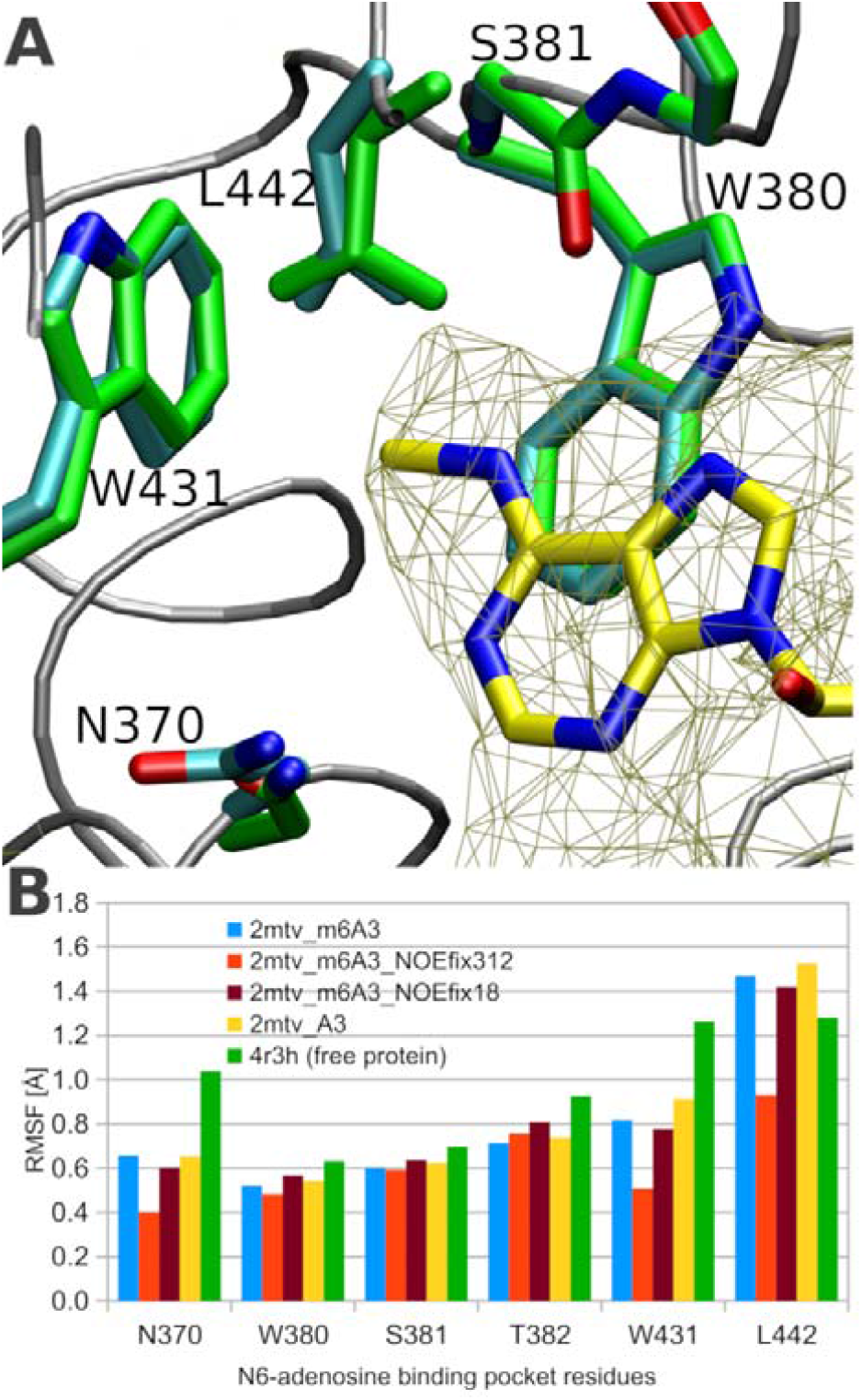
(**A**) Overlay of the average structures from 2mtv_m^6^A_3_ (protein/RNA complex; cyan carbons) and 4r3h (free protein; green carbons) simulations, respectively. The protein backbone and the bound m^6^A_3_ from the complex are shown in gray tube and with yellow carbons, respectively. In 4r3h simulations, the m^6^A_3_ binding pocket was composed and fully accessible for solvent molecules. The density grid of water molecules in the 4r3h simulations is indicated by the dark yellow mesh grid, overlapping with the m^6^A_3_ in the complex. (**B**) Graph of average per-residue root mean square fluctuation (RMSF) around the average simulation structure of selected systems, for residues lining the m^6^A_3_ binding pocket. With exception of L442, the range of fluctuations was greatest in simulations of the free protein.

### Methylamine probe mimics binding of the m^6^A to YTH

In light of the above results, we wished to directly explore the binding process of the m^6^A_3_ nucleotide within the YTH binding pocket and the associated solvent dynamics. However, in completely unbiased simulations, such process is unlikely to be observed on an affordable simulation timescale. For instance, in all our 2mtv_m^6^A_3_ simulations, we did not observe any signs of unbinding for the m^6^A_3_ nucleotide. A common approach in computational studies of substrate (un)binding is to induce or speed up the process by applying force potential bias along a single or multiple spatial coordinates, known as collective variables.^53^ However, results of such calculations can depend on the chosen collective variables, as they represent prior information (reaction coordinate) of the process with unavoidable dimensionality reduction.^16^ Here, the sampling of such process is further complicated by extensive involvement of the bulk water molecules. We faced such sampling limitations in our free-energy TI calculations investigating unbinding of the methyl group in the framework of the thermodynamic cycle (see Supporting Information). Thus, as an alternative approach to visualize accommodation of the methyl group in the binding pocket in the context of unbiased MD simulations, we have constructed a system in which we added twenty methylamine molecules into the water box surrounding the free YTH domain (Table 1). The methylamine is a structural mimic of the methylated amino group of m^6^A. In the subsequent simulations, we have observed methylamine spontaneously binding in the m^6^A_3_ binding pocket in ca 45% of simulation frames with maximum and average observed lifetimes of 203 ns and 68 ns, respectively. An essential observation was that long-term binding of a single methylamine in the m^6^A_3_ binding pocket on timescale of hundreds of nanoseconds would occur only in situations where the incoming methylamine would successfully displace all water molecules from nearby the W431 side-chain and assume a position very similar to the amino group of m^6^A (Supplementary Figure S3). This is consistent with a recent study which showed that release of a water molecule from this region is advantageous in terms of free energy.^18^ As expected, the methylamine was not completely selective for the m^6^A_3_ binding pocket and bound to YTH also at other locations (Supplementary Figure S4) although these were considerably less attractive, with the strongest alternative binding site having occupancy and maximum lifetime of 11% and 73 ns, respectively. Due to the imperfect selectivity of the methylamine, we cautiously refrain from making too strong conclusions from these simulations. However, they provided an important indication that displacement of water molecules from between the incoming nucleotide and the W431 side-chain is a necessary step for a long-term substrate binding. Such result is consistent with the active role of water molecules in binding of RNAs by proteins, as shown by earlier studies.^19, 54^-55

### G_2_ nucleotide is specifically recognized by a combination of direct H-bonding and water-mediated interactions

In experimental structures, the G_2_ is in *syn* and forms base stack with L383 side-chain and the G_2_(O6)/V385(N) H-bond.^13, 22^ Both the base orientation and its interactions were maintained in our simulations with reversible fluctuations. Simulations further predicted a G_2_(N1)/N386(OD1) H-bond (Figure 5A). All of these interactions are specific for guanosine and are the likely basis of the preferential recognition of 5’-Gm^6^A-3’ motifs by YTHDC1.^13, 22, 24^

**Figure 5.**
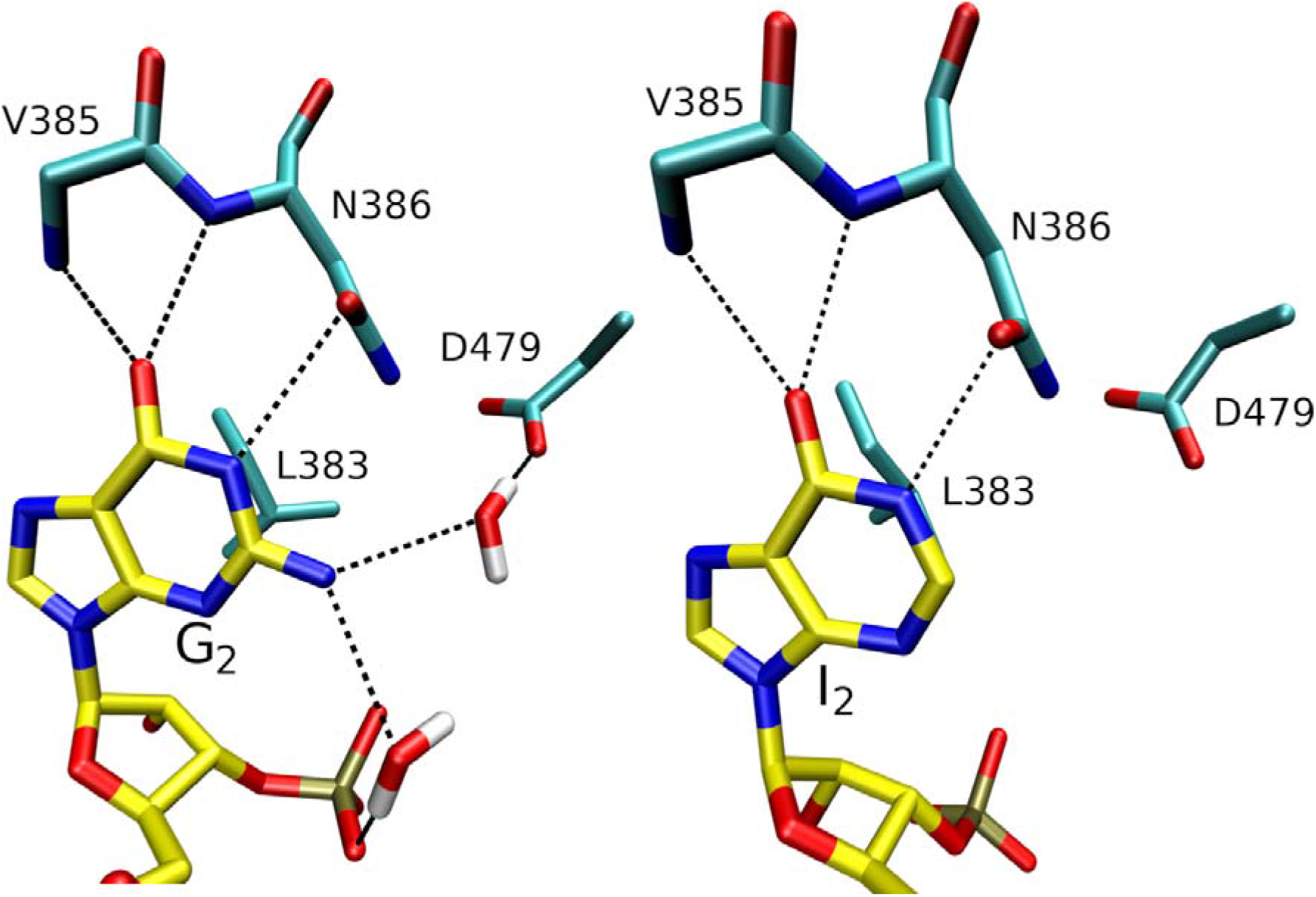
Protein/RNA binding pockets of G_2_ (left) and I_2_ (right) nucleotides in MD simulations of the YTH domain. Their binding was identical except for the missing water-bridge interactions in the I_2_ system.

In addition to the above, there was also a water bridge between G_2_(N2) and D479(OD) (Figure 5) formed in simulations. The same water bridge is observable also in the X-ray structure of YTHDC1 complex.^22^ It was yet another water-bridge formed between RNA and D479 side-chain, the latter of which also coordinates water molecules involved in m^6^A_3_ binding (see above and Figure 2). We subsequently performed simulations where we substituted G_2_ with inosine, thus abolishing the water-bridge between G_2_ and D479 by removing the 2-amino group. Surprisingly, elimination of this water-bridge had a broader influence on the network of water-mediated interactions in the region, including the water bridges within the m^6^A_3_ binding pocket. This was caused by the D479 shifting its position in presence of I_2_ (Figure 6). The significance of these changes for protein/RNA binding affinity is unclear. We would expect that the main contribution towards the preference for 5’-Gm^6^A-3’ motifs by YTHDC1 is rather supplied by the direct protein/RNA H-bond interactions, all of which remain the same for both G_2_ and I_2_. But it was previously shown by some of us that abolishment of intermolecular as well as intramolecular water bridges can lower the binding affinity of protein/RNA complexes in unexpected ways.^19^ Therefore, we further examined the effects of the G_2_→I_2_ substitution by performing TI free-energy calculations and NMR and ITC experiments. All three methods explicitly account for possible changes in the water network. A direct detection of the structured water molecules by NMR methods such as the CLEANEX-PM^56^ is unlikely to succeed given the relatively large flexibility of the region (Figure 4) and the known problems in interpreting the hydrogen exchange rates even under more ideal circumstances.^57^ However, NMR can detail the effects on the solute in terms of chemical shift perturbations (CSPs) and possibly elucidate the connection between the G_2_ and m^6^A_3_ pockets.

**Figure 6.**
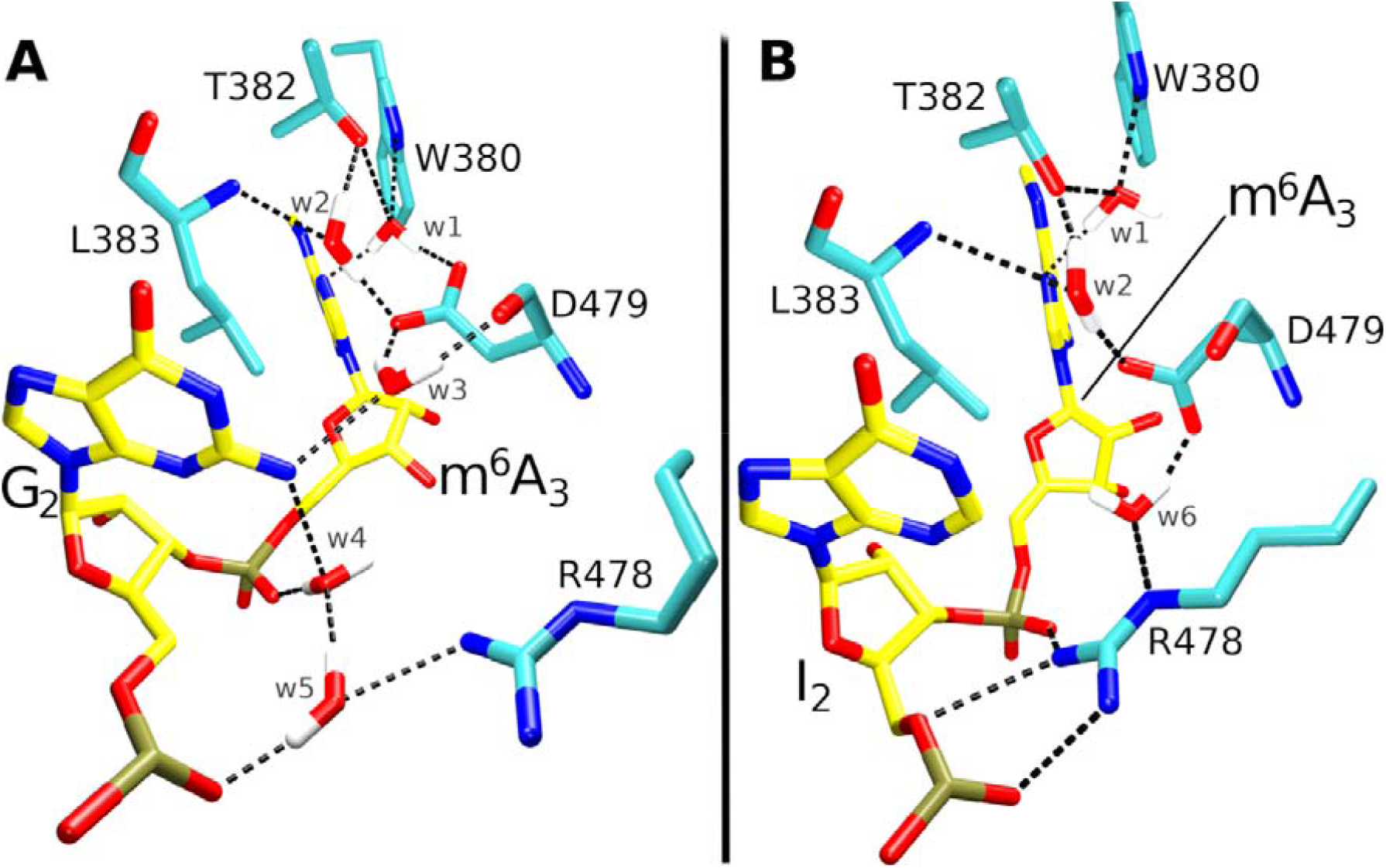
(**A**) Detailed view of the network of water-mediated interactions connecting the G_2_ and m^6^A_3_ binding pockets in MD simulations. (**B**) Absence of the N2 amino group in I_2_ disrupts the arrangement and causes changes to propagate across the entire network. The individual water bridges and protein/RNA interactions are labeled.

### Both calculations and experiments indicate a very minor reduction in binding affinity for the I_2_ RNA

We performed two independent TI calculations of the G_2_→I_2_ alchemical change that yielded ΔΔG values of 1.0 ± 1.3 and 1.5 ± 0.5 kcal/mol, respectively, for the relative stability of the two complexes. Although the trend is clear, we note the relatively large statistical uncertainty of both calculations, which, in case of the first calculation, exceeds the absolute value of the result. This may be caused by insufficient sampling of the water molecules which constitute the water bridges and directly interact with the atoms of the base undergoing alchemical transformation. For further details of the computational setup and results, see Supporting Information.

We also performed ITC measurements to experimentally determine the effect of the G_2_→I_2_ substitution on the binding affinity. The ITC yielded K_D_ values of 25 nM and 40 nM for the G_2_- and I_2_-containing substrates, respectively (Figure 7). Thus, the ITC also indicated a reduction of binding affinity, albeit merely by ∼0.3 kcal/mol. This supports our original assumption that preference for 5’-Gm^6^A-3’ motifs is primarily determined by the direct protein/RNA H-bonds with the G_2_ and not by the water-bridges.

**Figure 7.**
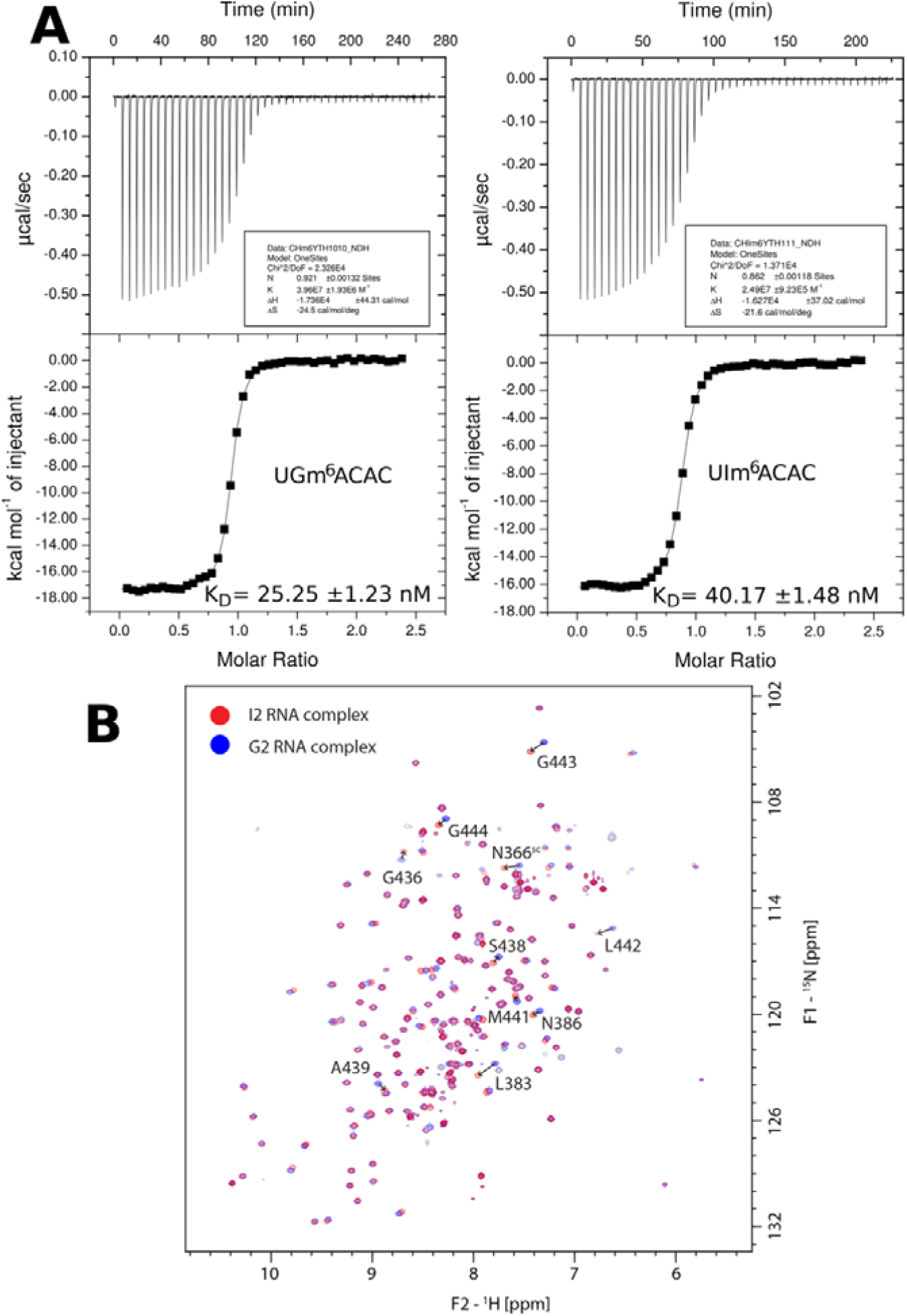
(**A**) ITC data recorded with the YTH domain of YTHDC1 bound to 5′-UGm^6^ACAC-3′ (left) or 5′-UIm^6^ACAC-3′ (right) RNAs. The K values are shown. Note that the earlier studies show variations in absolute K_D_s for the G_2_ substrate which we attribute to different compositions of the experimental buffers.^13, 22, 24^ (**B**) Superposition of ^15^N,^1^H-HSQC spectra of G_2_ and I_2_ protein/RNA complexes with 1:1 molecular ratio. Assignments of signals with largest chemical shift changes between the two complexes are shown and their directions indicated by arrows.

### NMR spectroscopy details structural effects of the G_2_→I_2_ substitution

The ^15^N,^1^H-HSQC spectra of the G_2_ protein/RNA complex measured for this work were highly similar to those measured earlier (Supplementary Figure S1).^13^ We attribute the small differences in peak positions to differences in NMR buffer compositions (see Methods). The I_2_ protein/RNA complex, measured for the first time in this work, also yielded very similar spectra (Figure 7) with only minor differences. Interestingly, the chemical shift changes were observed for the residues involved in the water interaction network between G_2_ and m^6^A_3_ as suggested by the simulations (Figure 6). For example, there was a minor downfield shift of 0.065 ppm for the W380(HE1) atom which is located deep in the m^6^A_3_ binding pocket and which is water-bridged with the D479(OD) in G_2_ but not in I_2_ simulations. A downfield shift of 0.170 ppm was also observed for the L383 backbone amide which, according to the simulations, is also involved in the aforementioned water network (Figure 6). Although the differences are very minor, the data confirms that there is a weak structural link between the G_2_ and m^6^A_3_ binding pockets sensitive to the I_2_ substitution. Therefore, even though this is not a direct proof of the water network, we suggest the observed chemical shift differences of the specific protein residues are consistent with its existence as suggested by the simulations.

Additional chemical shift differences were observed (Supplementary Figure S5), most of them modest and localized directly around the G_2_/I_2_ nucleotide binding site. Namely, there was a 0.065 ppm downfield shift of the ^1^H signal of N386 backbone amide which was predicted by simulations to interact with the O6 atom of both G_2_ and I_2_ (Figure 5). There was also a strong downfield shift of 0.155 ppm detected for one of the N366 side-chain amide signals (Figure 7). In simulations, this residue sometimes formed an H-bond with the m^6^A_3_ phosphate which in turn formed an intramolecular water bridge towards the N2 atom of G_2_ (Figure 6) as well as a water bridge with the R478 side-chain. In 2mtv_I_2_ simulations, these water bridges were abolished in favor of a direct interaction between the m^6^A_3_ phosphate and R478 (Figure 6). The remaining significant chemical shift differences were observed for the backbone amides of the β4/β5 protein loop (residues 436 to 446). This could be related to the L442 being one of the residues lining the m^6^A_3_ binding pocket (Figure 2). Lastly, we estimate the contribution of ring current intensities of hypoxanthine and guanine to the chemical shifts differences described above to be negligible.^58-59^ We base this on the small size of ring current shifts calculated for the G_2_ complex and the fact that the ring current intensities are only 8% larger for hypoxanthine compared to guanine.^60^

### D479A substitution destabilizes binding of the m^6^A_3_

The D479 side-chain formed a centerpiece of the water network connecting the G_2_ and m^6^A_3_ binding pockets (Figure 6) and so we decided to simulate YTH D479A mutant complex (Table 1) with the expectation that it will radically disrupt the water network. This was partially confirmed as the water bridges directly coordinated by D479 (Figure 6) were abolished in presence of A479 (Figure 8). The influence on the rest of the water molecules, not directly coordinated by D479, was more limited (Table 2). However, there was a curious effect on the w1 water bridge which directly participates in recognition of the m^6^A_3_ base (Figure 6). Namely, the water at this position would sometimes fail to properly exchange with the bulk solvent, leaving the site temporarily unoccupied and the many donors and acceptors engaged by this water unsatisfied. In other words, the occupancy of the w1 water bridge was significantly reduced by the D479A substitution (Table 2). In 2mtv_m^6^A_3_ simulations, the exchange of water molecules forming the w1 water bridge was always seamless, with the departing water molecule immediately replaced by another one from the coordination shell of the negatively charged D479 side-chain. Such mechanism of exchange is not available in presence of A479 and as a result, it could take time for new water to spontaneously enter the pocket after the previous w1 water is expelled in simulations. In the simulation frames where the w1 water bridge was temporarily absent, we sometimes observed reversible shift of the m^6^A_3_ base into the void left behind by the missing w1 water bridge. This shift was also associated with formation of an alternative “w1a” water bridge between T382(OG) and m^6^A_3_(O2′) atoms. In 2mtv_m^6^A_3_ simulations, both of these atoms were forming water-mediated and direct interactions, respectively, with D479 (Figure 2 and Figure 6). We suggest the D479A mutant system is attempting to compensate for the missing water bridges with D479 by shifting the m^6^A_3_ nucleotide into a position where favorable water bridges can be established once more. However, the shift also leads to loss of the vdW interaction between the N6-methyl group of m^6^A_3_ and the highly conserved W431.

**Figure 8.**
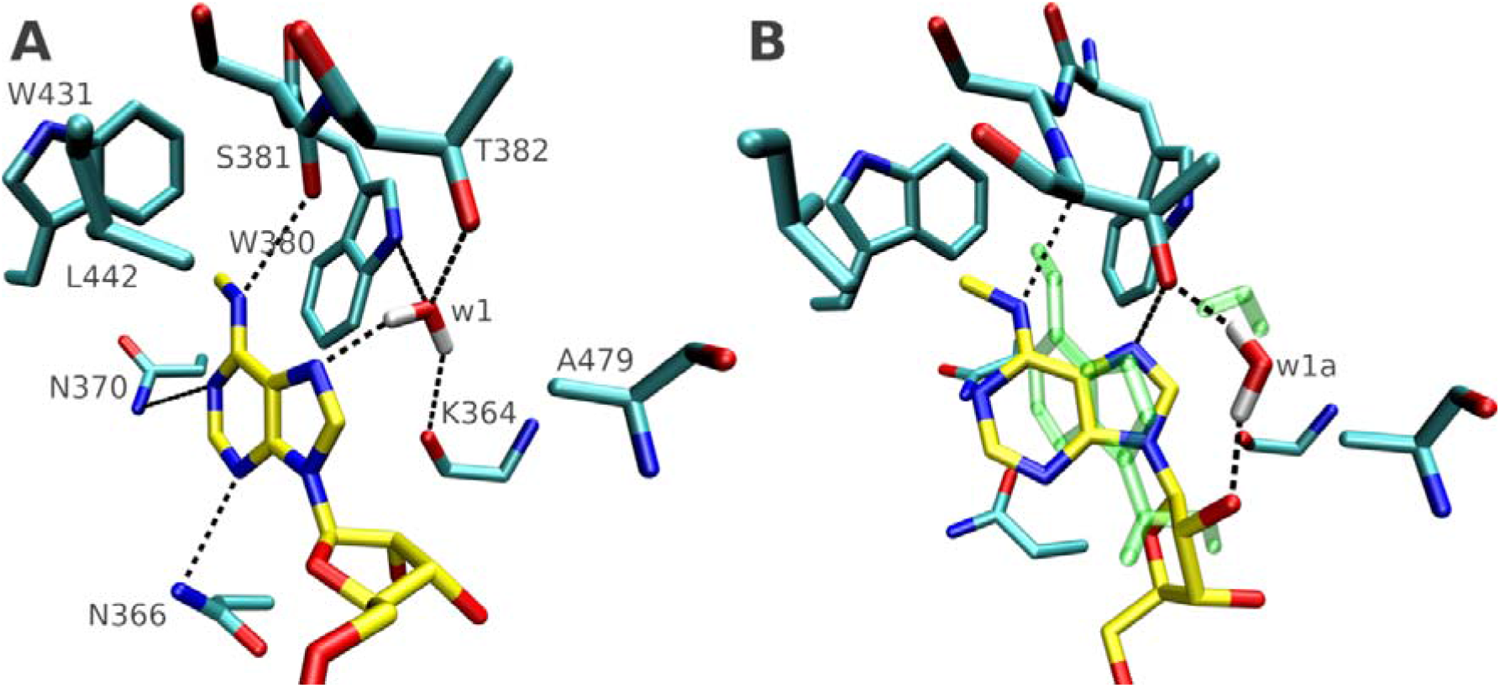
(**A**) Detail of the m^6^A_3_ binding pocket in 2mtv_D479A_m^6^A_3_ simulation when w1 water bridge was present. The amino acids forming the pocket and participating in the water bridge are labeled. (**B**) In some cases, the w1 water failed to exchange with the bulk which then caused the m^6^A_3_ nucleotide to move into the resulting void and form an alternative w1a water bridge. The shift disrupted many of the protein/RNA interactions native to the m^6^A_3_ binding pocket. The position of the m^6^A_3_ nucleotide and w1 water from panel A is shown as green transparent overlay for comparison.

In conclusion, simulations with the D479A substitution provide further evidence for the importance of the water network for establishing suitable arrangement of H-bond and vdW interactions ideal for m^6^A_3_ binding. The D479 side-chain facilitates seamless exchanges with the bulk solvent for the structured water molecules within the pocket while also establishing hydration patterns conducive for the m^6^A_3_ binding.

### Use of NOEfix function to further stabilize MD simulations of NMR structures

A common way to evaluate simulation force-field performance is comparison with the upper-bound NOE distances measured by NMR spectroscopy.^38, 61-65^ Here, we observed that over 90% of both intramolecular and intermolecular NOEs were satisfied in the standard 2mtv_m^6^A_3_ simulation ensemble (Table 3) which we consider very satisfactory for this type of biomolecule.^38^ Still, some of the NOE distance violations in our simulations occurred in vicinity of the G_2_ and m^6^A_3_ binding pockets (the primary interest of our study; see above). To verify our simulations and further improve the overall agreement with the experimental data, we formulated and tested the “NOEfix” potential, which is formally analogous to HBfix, gHBfix and tHBfix approaches tuning stability of H-bonds.^41, 66-67^ NOEfix introduces a mild stabilizing 1 kcal/mol energy bias between two hydrogens within interval of ld; d X 1.3J where *d* is experimental NOE upper bound (Figure 9A). Unlike traditional flat-well distance restraints commonly utilized for NMR structure refinement, the NOEfix does not produce any force potential for distances outside of the specified and limited distance range (Figure 9B).

**Table 3.** Percentage of satisfied NOE upper bound distances in MD simulations of the YTH domain complexed with 5′-C_1_G_2_m^6^A_3_C_4_A_5_C_6_-3′ RNA.

| simulated system | percentage of satisfied NOE distances |  |  |
| --- | --- | --- | --- |
|  | protein | RNA | protein/RNA |
| 2mtv_m<sup>6</sup>A<sub>3</sub> | 93% | 96% | 94% |
| 2mtv_m<sup>6</sup>A<sub>3</sub>_NOEfix312 | 96% | 98% | 99% |
| 2mtv_m<sup>6</sup>A<sub>3</sub>_NOEfix18 | 94% | 98% | 99% |

**Figure 9.**
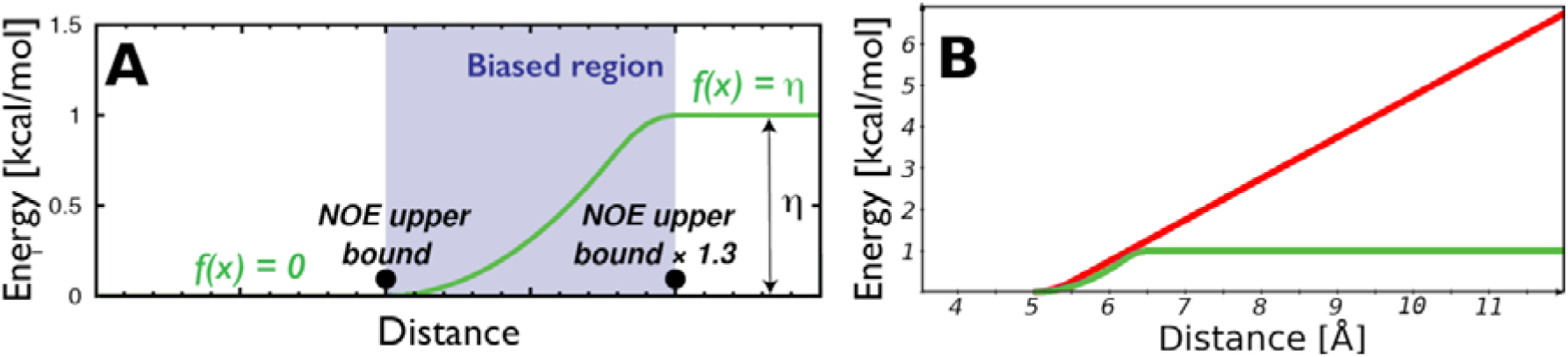
(**A**) The NOEfix potential (green curve) produces non-zero forces only in a narrow region between the NOE upper bound distance and the same distance multiplied by factor of 1.3. A smoothly increasing potential is applied in this range until it reaches the maximum energy penalty η. In this fashion, a steeper NOEfix potential is utilized for stronger (shorter) NOEs while a more gentle potential is present for the weaker (longer) NOEs. The distance range and η values are adjustable. For details of the technical implementation of the potential function, see Refs.^41, 66-67^. (B) Comparison of standard restraint with force constant of 1 kcal/mol/Å^2^ (red curve) and NOEfix (green curve) applied to NOE with upper bound of 5 Å. Much stronger force constants are typically utilized with standard restraints. NOEfix is a pragmatic means of introducing some bias based on the primary NMR data to compensate for the force-field deficiencies and to verify the performance of fully unbiased simulations while maintaining much greater possibility for conformational exploration than what would be observed with traditional restraints where the energy penalty keeps increasing with distance.

In our initial test, we applied NOEfix for all the violated NOE distances observed in 2mtv_m^6^A_3_ ensemble (considering all eight simulations together), corresponding to 312 NOE distances. In this 2mtv_m^6^A_3__NOEfix312 simulation ensemble, we observed significant improvement in the number of both intra- and intermolecular satisfied NOEs in simulations (Table 3). However, use of this many NOEfixes noticeably reduced the fluctuation range of the residues lining the m^6^A_3_ binding pocket (Figure 4) and some of the NOEfixes were never violated at any point of the simulations (Supplementary Figure S6). This goes against the original idea of including experimental information in the simulation ensemble without excessively hindering the ability to explore conformational space. In other words, such indiscriminate use of NOEfix has likely a similar effect as mere restraining; see Supporting Information for more details.

In our second test, we limited application of NOEfixes for the violated NOE distances occurring for residues within 6 Å of the bound RNA. Only the single largest NOE distance violation per each protein residue was considered, obtaining a selection of 18 NOEfixes (see Supporting Information for the complete list). In this 2mtv_m^6^A_3__NOEfix18 ensemble, we observed improvement in the intermolecular satisfied NOEs that was on par with the 2mtv_m^6^A_3__NOEfix312 ensemble despite significantly fewer NOEfixes being used. Overall, we observed 64 fewer NOE violations in 2mtv_m^6^A_3__NOEfix18 compared to 2mtv_m^6^A_3_, though 35 new NOE violations appeared. Importantly, use of NOEfix lowered number of NOE distance violations around the G_2_ and m^6^A_3_ binding pockets (Supplementary Figure S7) in both 2mtv_m^6^A_3__NOEfix312 and 2mtv_m^6^A_3__NOEfix18 simulation ensembles. Almost the same improvement could be achieved with 18 NOEfixes while the use of fewer, carefully selected NOEfixes does not repress the dynamics (Figure 4 and Supplementary Figures S8 and S9). We note the NOEfix ensembles provided very similar structural information as the 2mtv_m^6^A_3_ ensemble (Figure 4 and Figure 6), demonstrating that the main results reported in this work are not affected by force-field related deviations from the primary experimental data. The YTH is a well determined system with reasonable force-field performance. Possible uses and advantages of NOEfix with less well-behaved systems are stated in the Discussion.

## Discussion

### Structural hydration plays a key role in recognition of the N6-methylated adenosine by the YTH domain and in discrimination against unmethylated adenosine

Our simulations provided unique insights into interrelation between hydration dynamics and protein/RNA recognition by revealing a set of water bridges in the vicinity of the m^6^A_3_ binding pocket. A key role of water molecules for the m^6^A_3_ binding and discrimination against A_3_ was further shown by our simulations with the unmethylated A_3_ nucleotide. In the bound state, the A_3_ nucleotide had two dynamically competing binding states which both resembled the ground state of m^6^A_3_ when considering the position of the base. In the dominant state (∼90% occupancy), there was no water molecule inside the pocket. This left an unsaturated H-bond donor as well as not fully saturated vdW surface within the pocket due to absence of the N6-methyl group on A_3_. In other words, in the dominant binding state, the protein could not fully optimize its interactions with the unmethylated adenosine. In our simulations, the system consequently sampled an additional auxiliary binding state (∼9% of time) where a single water molecule transiently invaded the binding pocket in the A_3_ simulations (Figure 3). This water molecule occupied the space in which the N6-methyl group would normally be located for the m^6^A_3_ substrate, forming a single H-bond with the H61 hydrogen of A_3_, as well as compensating for the loss of the methyl-group vdW contacts (Figure 3). However, this single water molecule was unable to establish any other H-bonding interactions. Such an arrangement likely carries a large free-energy penalty since a single water molecule can potentially form up to four H-bonds.^68^ Furthermore, buried water molecules are entropically unfavorable.^69^ Thus, for distinct structural reasons, the A_3_ molecular interactions with YTH were suboptimal both in the dominant and auxiliary binding states.

What is striking is that the auxiliary binding state appears to also be a transition (or intermediate) state along the coordinate corresponding to the (un)binding process for A_3_. Specifically, our simulations showed that the invading water molecule associated with the auxiliary state was either expelled again from the pocket after some time, thus reverting to the dominant binding state, or it attracted additional water molecules from the bulk which, then, gradually displaced the A_3_ nucleotide from the pocket (Figure 3). Based on this data, we suggest that binding of m^6^A_3_ and A_3_ substrates by the YTH domain differs not only in the free energy of the final states, as shown by the experiments,^13, 22^ but that the kinetics and pathways of the two bindings could also be different. Namely, the recognition of the m^6^A_3_ and A_3_ by the YTH could correspond to two-state and four-state bindings, respectively (Figure 10). This could have biological implications as the dynamics of the single water molecule observed in the auxiliary binding state of unmethylated A_3_ and its ability to attract even more waters from the bulk could aid in more frequent displacements of the unmethylated adenosine from the pocket. In the presence of other RNA molecules, it could lead to a faster substrate cycling, and, in turn, allow the YTH domain to more quickly locate the N6-methylated adenosine among a pool of cellular RNAs where unmethylated adenosine can be expected to be much more prevalent. Incompatibility between the A_3_ binding and relaxed water network is assumed to substantially increase “local” k_off_ kinetic constant of the nucleotide.

**Figure 10.**
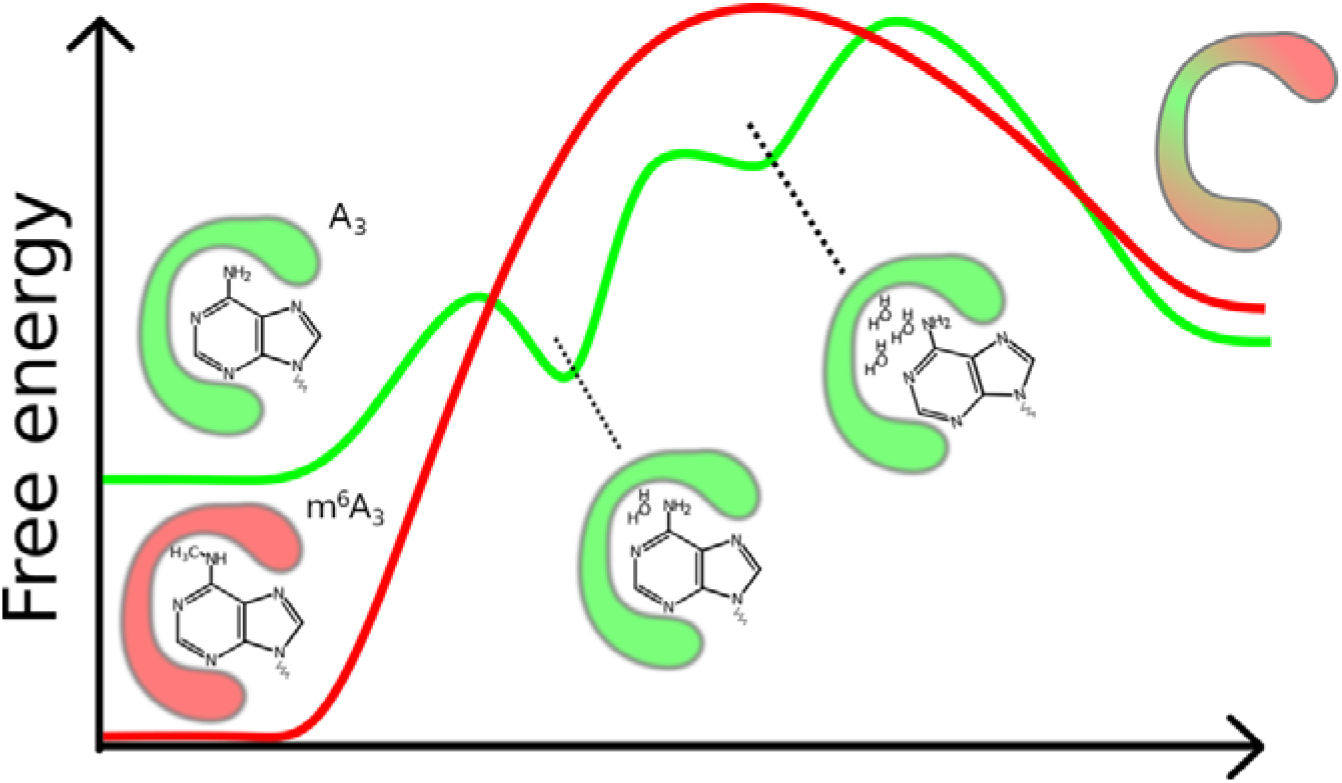
Scheme of the suggested differences in binding states of the N6-methylated (red) and unmethylated (green) adenosine by the YTH domain of YTHDC1. The energy curves are merely illustrative of simulation trends and are not a result of measurements or free-energy computations.

Lastly, we note that our simulations of the free YTH domain showed that the binding pocket is preformed and fully accessible to the solvent; in other words, it is hydrated at all times (Figure 3). Thus, the substrate must displace these water molecules during the binding. The simulations generally indicate that there are several mechanisms which contribute to the preferred binding of the N6-methyl adenine over adenine, albeit they do not allow an exact decomposition of the process. Our thermodynamic integration calculations revealed that the free energy of solvation is ∼0.7 kcal/mol higher for adenine than for the N6-methyl adenine. The desolvation of the Watson-Crick edge of both bases during the binding could then be one of the contributing factors to the YTHDC1’s substrate preferences. In other words, the A_3_ is more attractive for waters and consequently less able to squeeze them out of the binding pocket, potentially resulting in lower affinity towards YTHDC1 compared to m^6^A_3_. The importance of displacing the water molecules from the pocket was demonstrated by simulations of the free protein in presence of methylamine (a molecular mimic of the N6-methylated amino group) which was able to (meta)stably bind to the m^6^A_3_ binding pocket only after excluding all water molecules from the pocket (Supplementary Figures S3 and S4). Importantly, position of the bound methylamine confirms that the pocket is perfectly evolutionarily optimized to recognize the modified nucleotide. From the qualitative point of view of energy or free-energy landscapes,^70-71^ one could also argue that the simulations nicely illustrate how in absence of the N6-methyl group the interaction interface becomes considerably more frustrated upon binding. Release of the water molecules from binding interfaces is generally known to promote folding and binding by increase of the solvent entropy (the existence of the preformed pocket together with the smooth binding of methylamine probe actually locally resembles the idealized “lock and key” binding mechanism which was suggested to lead to large solvent entropy stabilization.^71^. The key difference between m^6^A_3_ and A_3_ is a clear frustration of the interactions (and thus unoptimized enthalpy stabilization) when the latter moiety enters the pocket, irrespective of whether the water molecules are completely displaced or one water molecule remains temporarily trapped.

We acknowledge the simulations are likely not long enough to capture all the possible structural micro-arrangements. Nevertheless, they provide an illuminating view into the role of water dynamics during binding/unbinding of a N6-methylated adenosine in the binding pocket of YTH and how this can be used for discrimination against an unmethylated substrate. We propose that similar mechanisms might be used by some other proteins which recognize residue methylation through similarly structured binding pockets.^8,11^

### The YTH domain can tolerate inosine as the nucleotide preceding m^6^A_3_

Earlier experiments have shown that the YTHDC1 binds the RNA substrate more strongly when guanosine is the nucleotide immediately preceding the m^6^A (the G_2_ in our work).^13, 15, 22^ Our simulations showed that the G_2_ formed several specific albeit fluctuating interactions with the YTH. Among these was a water bridge interaction between the N2 amino group and the D479 side-chain (Figure 6). We performed several simulations in which we mutated the G_2_ into inosine to abolish this water bridge. We subsequently quantified its effect on RNA binding affinity by performing TI calculations and ITC measurements (Figure 7A), which indicated a minor free-energy penalty for the I_2_ complex of 1.0-1.5 kcal/mol and 0.3 kcal/mol, respectively. Furthermore, NMR spectroscopy identified residues affected by this modification and suggested that the absence of the N2 amino group in the I_2_ RNA complex and the loss of the water bridge are partially compensated for by strengthening of some of the other protein/RNA interactions. Our work thus suggests that the YTH domain of YTHDC1 can recognize both 5′-Gm^6^A-3′ and 5′-Im^6^A-3′ motifs with only a minor free-energy penalty for the latter.

### Water bridges facilitate communication between the G_2_ and m^6^A_3_ binding pockets

The simulations revealed a network of water bridges, connecting the G_2_ nucleotide through its 2-amino group with residues lining the m^6^A_3_ binding pocket (Figure 6). Lifetimes of the individual waters in this network were on timescale of tens to hundreds of nanoseconds (Table 2) while common hydrations sites around biomolecules have sub-ns time scale.^16^ We showed that the water network could be altered by G_2_→I_2_ substitution which abolishes water bridges made by the 2-amino group (Figure 6), although the direct H-bonds with the protein are not affected. Chemical shift changes were detected by NMR for the protein residues deep within the m^6^A_3_ binding pocket, giving an indirect experimental support for existence of the water network (see above and Figure 7B). We further explored the intricacies of the water network in our simulations of the YTH protein/RNA complex with D479A mutation. The side-chain of D479 is specifically involved in coordinating the water network (Figure 6) and its removal had profound effects. Namely, without D479, the single water bridge deep in the m^6^A_3_ binding pocket was sometimes unable to make a complete exchange of water with the bulk solvent, leaving the hydration site temporarily vacant. In some simulations, this compromised the m^6^A_3_ binding (Figure 8). This illustrates the importance of the water molecules both for establishing the specificity of the N6-methylation recognition (see above and Figure 3) as well as proper arrangement of the binding pocket.

### Agreement between MD simulations and experimental data can be improved and validated with NOEfix

The simulations showed a reasonable force-field performance in describing parts of the protein/RNA interface related to recognition of the G_2_ and m^6^A_3_ nucleotides (the primary focus of this study). Nevertheless, there were some NOE distance violations reported for this region in the standard simulations. These violations could potentially affect the results but applying traditional NMR restraints would have severely limited the dynamics. Therefore, we instead applied the newly-developed “NOEfix” function (Figure 9). Based on the same underlying math as the HBfix,^16, 41, 66^ the NOEfix increases the agreement between the simulation and the experimentally measured upper-bound NOE distances (Table 3). In the two simulation tests, we applied the NOEfix to either all observed NOE distance violations (312 NOEfixes), or to selected NOE distance violations occurring close to the protein/RNA interface (18 NOEfixes). The latter choice is more appropriate as using too many NOEfixes at the same time can repress the structural dynamics (Figure 4 and Supplementary Figure S10). In both tests, the use of NOEfix corrected majority of the violated NOE distances but also introduced some new violations. This shows that simultaneous satisfactory description of all the NOE distances indicated by the experiment is not favored by the force field. While not fully successful, it is important to note that a perfect agreement with NMR data should not be realistically expected due to inherent errors of the experimental technique as well as the unphysical approximations of the molecular mechanical description. The NMR data used for comparison is also ensemble- and time-averaged^21^ and so the inability to satisfy all the NOE distances in simulations could also reflect insufficient sampling. The appearance of new NOE violations could also be a sign of an already excessive bias even when using only 18 NOEfixes. Possible improvements could be achieved by reducing the number of utilized NOEfixes even further. The system studied in this work is also well-determined, possessing very high density of NMR signals, many of which are structurally coupled.^13^ The NOEfix could be of greater benefit in systems where insufficient number of signals was detected due to, e.g. partial disorder or conformational transitions.

Obviously, the NOEfix approach does not represent a direct rigorous integration of the primary NMR data into the MD simulation, such as for example the maximum entropy approaches.^72-74^ However, it is easy and flexible in its implementation and can improve quality of the simulated ensembles even in cases when direct integration of the experimental data into simulation protocols is inconvenient due to, e.g., insufficient amount of the primary experimental data or large size of the system. Note that although the presently studied system provides a relatively large amount of primary experimental data compared to many other protein-RNA complexes, it could still be very challenging for a reliable unambiguous direct integration of experimental data with the simulations.

NOEfix could be particularly useful for resolving distance violations arising from minor force-field inaccuracies across multiple terms whose direct tuning would lead to problems in other parts of the system. Similarly as with the maximum entropy approaches, the NOEfix approach is motivated to minimize the effect on the simulation ensemble compared to the common restraints. However, NOEfix obviously affects the ensemble, as it primarily aims to reduce populations of all substates where the NOE distances are violated. This corresponds to an assumption that these substates are excessively populated due to force-field inaccuracies. The presently suggested NOEfix parameters were chosen rather intuitively and are not based on any rigorous approach. We plan to test applicability of NOEfix in the near future for a wide range of protein-RNA complexes, including those which are less well determined and show larger instabilities in the simulations.

### Concluding remarks

We have investigated dynamics of the YTH domain of the YTHDC1 protein and its binding to N6-methyladenosine containing RNA. We show that the mechanism of recognition, in addition to stabilization via the vdW interactions with the methyl group, includes contributions from structured water molecules. While long-residency water molecules are helping to cement binding of the base, unmodified adenosine is unable to keep water molecules from within the N6-methyladenosine binding pocket. Our study therefore provides an additional structural basis for the ∼50× reduction in affinity of unmodified adenosine containing RNAs compared to those harboring m^6^A. Water molecules are also involved in recognition of the G_2_ residue and there is an intricate network of water bridges connecting the two binding pockets. We used NMR to measure the effects of disrupting this network.

## Supporting Information

The following files are available free of charge https://pubs.acs.org. Additional details on simulation methods and results. Supplementary Figures.

## Acknowledgement

This work was supported by the Czech Science Foundation [grant number 20-16554S to M.K. and J.S]; NCCR RNA and Disease from the Swiss National Science Foundation to F.A.; and CESNET storage facilities [grant number CESNET LM2018140].

## TOC Graphic

