## Supplementary for "Recognition of N6-methyladenosine by the YTHDC1 YTH domain studied by molecular dynamics and NMR spectroscopy: The role of hydration"

### Supplementary Data

**Comments on starting structure selection for simulations of the protein/RNA complex.** The two structures of the YTH domain of YTHDC1 complexed with RNA available in the database (PDB: 2mtv, NMR structure, first conformer, and PDB: 4r3i, X-ray structure) show a highly similar protein/RNA interface.^1-2^ At the initial stages of the project, we have considered both structures as viable candidates for our MD simulations. However, since a significant portion of our study dealt with the use of NMR primary experimental data to stabilize the simulations (NOEfix function; see the main text) and the structures are very similar, we decided to use the NMR structure as start of our simulations. We stress that despite ultimately proceeding with the NMR structure, we consider the quality of the X-ray structure to be excellent and on par with the NMR structure. Where relevant, we utilize the X-ray structure as an additional reference in interpreting the simulation results.

Another reason for our decision was that for larger molecules such as protein-RNA complexes the standard equilibration protocol is not always fully efficient. Even essentially correct experimental structures can sometimes lead to excessive perturbation in initial stages of the simulations, since the starting Cartesian-coordinate configuration taken from the experimental structure is not relaxed with respect to the force field. The simulated molecule thus needs to dissipate the excess energy which can occur in the early stages of the production simulations. It may then occasionally lead to perturbations of the simulated structures.^3-4^ Parts of the YTH system were susceptible to these problems (see below). The possibility of using the NMR restraints in the initial portions of our simulations as part of our computational protocol (see the main text Methods) to mitigate such developments was then advantageous with the NMR structure.

**Changes of the protein/RNA interface related to the C_4_, A_5_, and C_6_ nucleotides outside the primary binding sites in preliminary simulations; use of Hbfix potential.**

Our preliminary set of MD simulations revealed perturbations of the native protein/RNA interactions formed between positively charged arginine and lysine side-chains and the phosphates of C_4_, A_5_, and C_6_ nucleotides. When occurring individually, these perturbations were reversible on our simulation timescale and likely represented realistic dynamics for the system and this type of interaction. In fact, such behavior has been previously reported for interactions formed by positively charged amino-acids and RNA phosphates.^5^ However, when these perturbations occurred simultaneously, they would become irreversible because the RNA would form various non-native intramolecular interactions. These non-native RNA self-interactions were very stable on our simulation timescale and essentially “locked” the RNA into a conformation in which the native protein/RNA interactions could no longer be reformed (Figure S11). This resulted into a loss of the protein/RNA interface for the C_4_, A_5_, and C_6_ nucleotides, but not the G_2_ and m^6^A_3_ nucleotides. We strongly suspect that the loss of the protein/RNA interface for the C_4_, A_5_, and C_6_ nucleotides and especially formation of intramolecular RNA interactions is a force-field error rather than indication of genuine biomolecular dynamics. It is highly consistent with various issues reported for simulations of single-stranded RNAs.^3^^,^ ^6-9^ It did not seem to affect the G_2_, I_2_, m^6^A_3_ and A_3_ binding sites significantly, on our simulation timescale. However, for sake of caution, we refrained from basing our conclusions (see main text) on such simulations. To obtain a more reliable simulation ensemble, we applied mild (1 kcal/mol in terms of potential energy) structure-specific HBfix potential^10^ to increase stability of two H-bonds natively formed by nucleotides A_5_ and C_6_ (A_5_(OP1)/K364(NZ) and C_6_(OP1)/K475(NZ); see main text Table 1). This choice was aimed at stabilizing the native interactions formed by the 3’-terminal parts of the RNA chain where the perturbations were seen earliest and most frequently in the simulations. At the same time, the use of mild HBfix far away from the G_2_ and m^6^A_3_ binding sites can be, in our opinion, safely assumed not to alter the dynamics for these binding pockets. This assumption is supported by the highly similar results of the simulations utilizing the NOEfix, where the aforementioned HBfix potentials related to the A_5_ and C_6_ nucleotides were not used. With the application of either the HBfix or the NOEfix, the spurious intramolecular RNA interactions (Figure S11) were not observed in the simulations (main text).

We also note that the spurious intramolecular RNA interactions were also observed in preliminary simulations performed with modified vdW parameters of phosphates and the OPC water model (bsc0χ_OL3__CP(OPC) force-field combination; see main text Table 1).^11-13^ These simulations were primarily used to verify that the same hydration sites would be observed with OPC as well as the SPC/E water molecules (see the main text).

**Methodological details of the thermodynamic integration (TI) calculations.**

We used soft-core potentials in our TI calculations. Using this method, the appearing and disappearing atoms (i.e. atoms unique to the initial or the final state) can be present in the system at the same time. Their vdW interactions with the rest of the system are smoothly switched off by utilizing a modified version of the vdW equation.^14^ To handle the electrostatic change, we used the traditional three-step methodology of discharging–transforming–recharging the mutated atoms.^15^ For one specific alchemical transformation, we also used the soft-core electrostatics method which allows alchemical transformation of electrostatic and vdW components of the atoms in a single step.^16^ We collected the derivatives of the force potential for each lambda window (*δV/δλ* values) at every integration step. We used the nine-point Gaussian quadrature to numerically estimate the final energy integral for each step using in-house scripts. We used a batch method to express the standard deviation of the calculations where each lambda window was divided into clusters of one million lambda values and their averages computed. The standard deviation was then derived from these averages.^17^ At the end, the final free-energy difference of the mutated protein/RNA complex (ΔΔG) was computed according to the thermodynamic cycle equation ΔΔG = ΔG_co_ – ΔG_s_ = ΔG_WT_ – ΔG_mut_. ΔG_co_ and ΔG_s_ represent the results of the alchemical TI calculations in the protein/RNA complex and in the single-stranded RNA, respectively. ΔG_WT_ and ΔG_mut_ are the dissociation energies of the wild-type and the mutated complex, respectively, as commonly measured in experimental setting. Due to validity of the thermodynamic cycle, the two approaches (experiment and computation) obtain a physically equivalent result (ΔΔG) within the approximation of the force field and sampling.^3^ No positional restraints were used in our TI calculations.

**TI calculations of the solvation free energy of the bases**. We constructed isolated A, m^6^A, and G bases (guanine was used for control) by removing the phosphate and sugar atoms from the nucleotides, retaining only the base and the C1’ and H1’ atoms. Two additional hydrogens were attached to the C1’ atom to form a methyl group cap for the N9 atom and the partial charges of the three hydrogens were manually adjusted to obtain a zero net charge of the system. The constructed systems were surrounded by a water box and then equilibrated (see the main text). We then performed five-point Gaussian quadrature thermodynamic integration calculations^15^ with each lambda window simulated for 20 ns in which we calculated the free energy of solvation for all three bases by gradually decoupling the nonbonded interactions of the base with the water box. Soft-core atoms were utilized to describe both the vdW and electrostatic changes. The calculated free energy of solvation was -12.53 ± 0.08, -11.86 ± 0.08, and -22.48 ± 0.09 kcal/mol for A, m^6^A, and G, respectively. The error of the calculations was estimated by the same method as for the other TI calculations (see above). The values calculated for A and G are in agreement with diverse earlier calculations^18-19^ which suggests that our procedure should correctly reflect the difference between adenine and N6-methyl adenine.

**Details of the thermodynamic integration calculations of the m^6^A_3_↔A_3_.**

Binding of methylated adenine in the YTH pocket is at first sight an ideal system for accurate alchemical free-energy computations such as the TI method (for a review discussing applicability of different free-energy computations for protein-RNA systems see Ref.^3^). Thus, in the first set of TI calculations, we examined the effects of the RNA binding containing either m^6^A_3_ or A_3_ on the complex stability. The experimental data indicated a ~50 fold reduction of binding affinity of the YTH domain for the standard adenosine-containing RNA substrate which corresponds to a free-energy penalty of ca 2.3 kcal/mol.^2^ Nevertheless, we wished to evaluate the quantitative effects of this modification also in the context of our simulations. To do this, we have performed both forward and backward TI calculations of the m^6^A_3_↔A_3_ substitution in the context of the protein/RNA complex and of the single-stranded RNA, using the standard thermodynamic cycle (see the main text Methods and above).^3^ Performing TI calculations in both directions seemed necessary since standard MD simulations indicated that one of the main differences between the two states were regular intrusions of a single or multiple water molecules into the m^6^A_3_/A_3_ binding pocket (see main text Figure 3) when standard adenosine was present. The backward calculation was performed with a single water molecule already present within the binding pocket. This water was seamlessly displaced by the growing N6-methyl group in the TI calculations in the lambda windows close to the m^6^A_3_ end-state. We expect that it may be challenging to sample such differences with the standard TI calculation setup without including the invading water molecule(s) into the list of appearing and disappearing atoms. However, inclusion of the water molecules would complicate the thermodynamic cycle and introduce a large source of error into the calculation. Since the invading water can exchange with bulk during the simulations, it would also necessitate the use of restraints to keep the alchemical water from exchanging with the waters from the bulk solvent. However, MD simulations show that the water is not present inside the pocket at all times (see main text), so that such a representation would also not be ideal. In other words, there is no single starting geometry which would be ideal for describing the m^6^A_3_↔A_3_ alchemical change in the context of TI calculations. For results of the m^6^A_3_↔A_3 ­­_TI calculations, see the main text.

**Details of the 2mtv_m^6^A_3__NOEfix312 simulations.**

The 221 NOE violations which were observed in the 2mtv_m^6^A_3_ ensemble were corrected in 2mtv_m^6^A_3__NOEfix312 ensemble. Of the remaining violations, 91 were observed in both ensembles and 79 violations were newly observed in the 2mtv_m^6^A_3__NOEfix312. Importantly, the absolute values of the remaining violations were significantly lowered compared to the standard ensemble (Figure S8). Still, the appearance of new violations suggests that using 312 NOEfixes is excessive. Also, the thermal fluctuations of some of the residues lining the m^6^A_3_ binding pocket were notably reduced in the 2mtv_m^6^A_3__NOEfix312 ensemble (main text Figure 4) and some of the NOE upper bound distances for which we applied NOEfix were not exceeded at any point of the simulation (Figure S6). Likewise, the potential energy associated with NOEfix quickly converged and remained close to constant in these simulations (Figure S10).

### Supplementary Figures

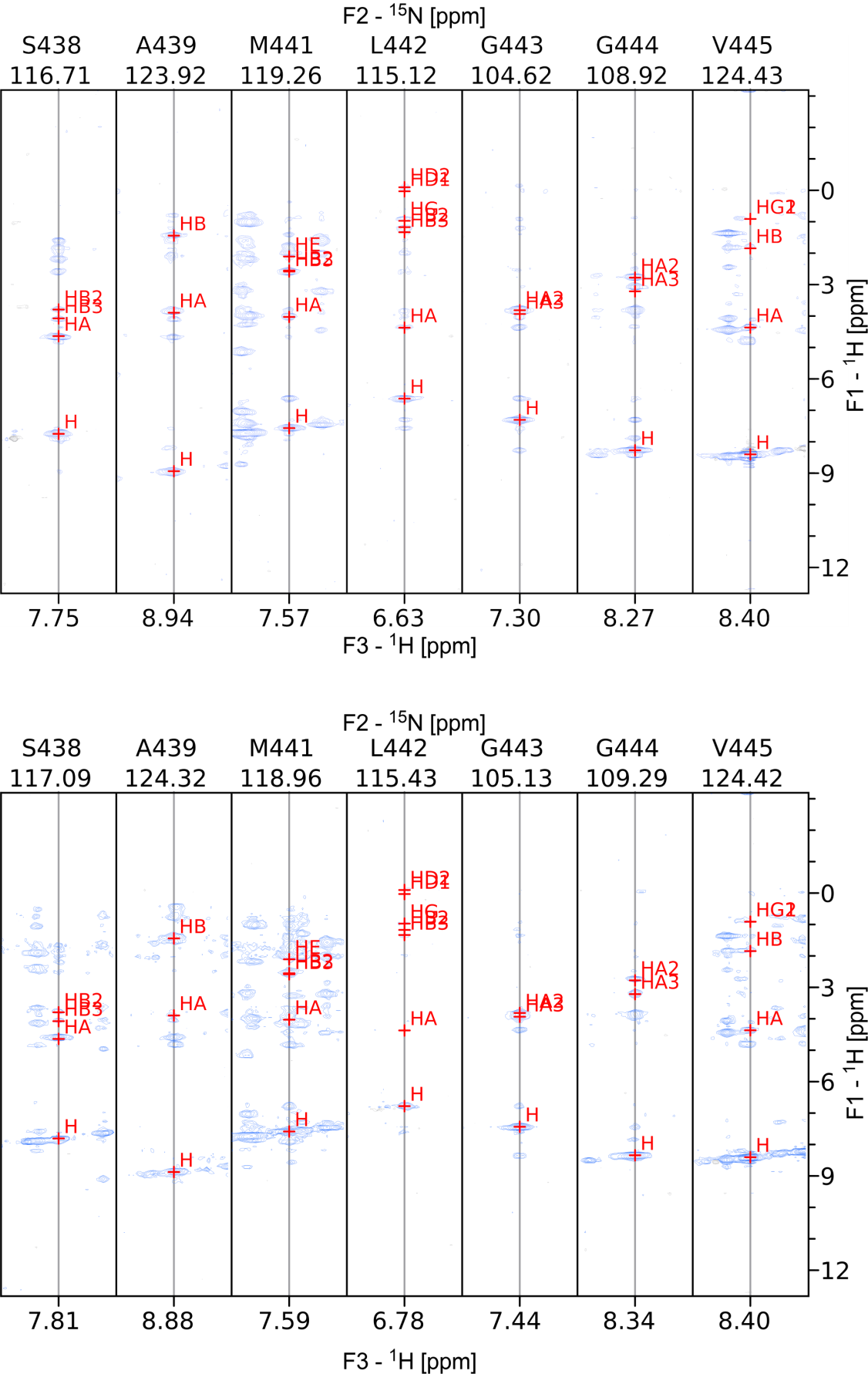

Figure S1. Comparison of the NOESY towers for residues 438-445 (labeled at the top of each strip) from the 3D ^15^N-resolved NOESY-HSQC of the G_2_ (top) and I_2_ (bottom) protein/RNA complexes. The high similarity allowed identification of the amide N-H positions in the ^15^N,^1^H-HSQC of the I_2_ RNA. The 438-445 residues had the largest combined chemical shift changes between the two structures. For reference, the ^1^H assignments are shown in red at the NOE positions in each strip.

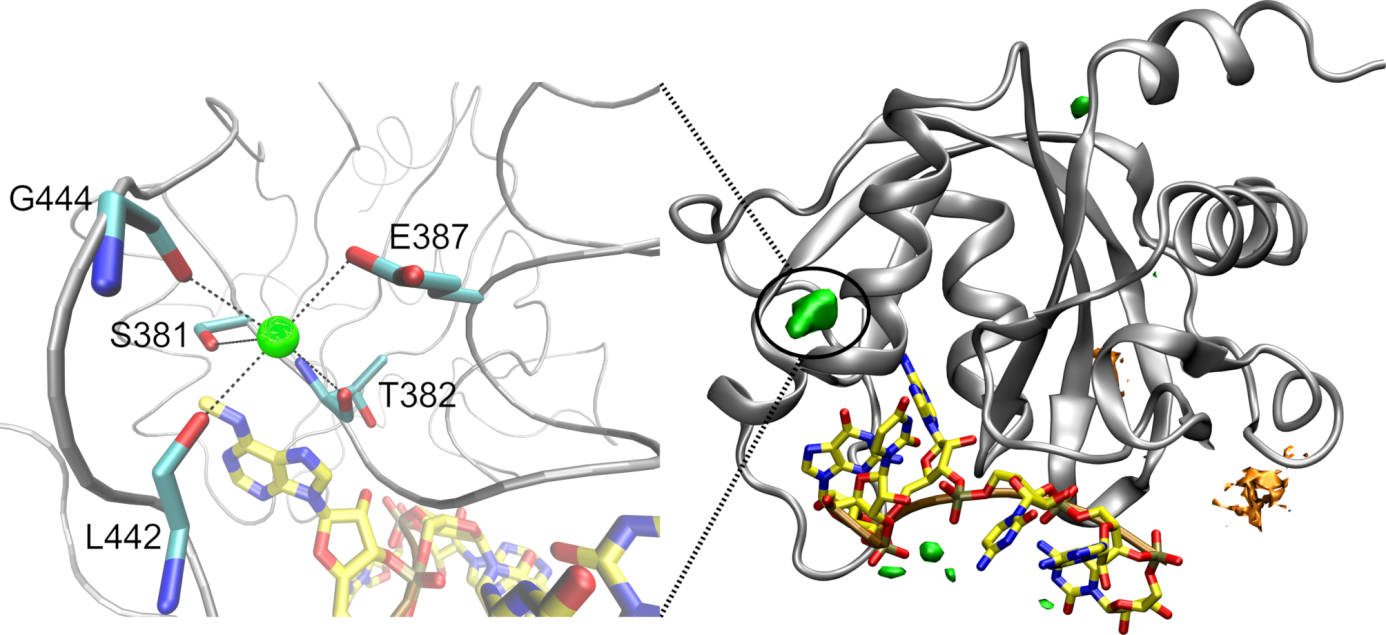

Figure S2. The most prominent potassium (green sphere) binding site in MD simulations (left) was observed near the G444(O), L442(O), T382(O), S381(OG), and E387(OE) atoms with an average occupancy in simulations of ca 25%. The coordination of the ion is indicated by black dashed lines and the coordinating residues are labeled. On the right are shown grid density distributions of potassium (green) and chloride (orange) ions in the combined ensemble of the 2mtv_m^6^A_3__NOEfix18 simulations. At least five points with highest occupancies are shown for each ion. The black circle indicates position of the potassium binding site shown in the left panel. The analysis reveals absence of any salient ion binding in the system.

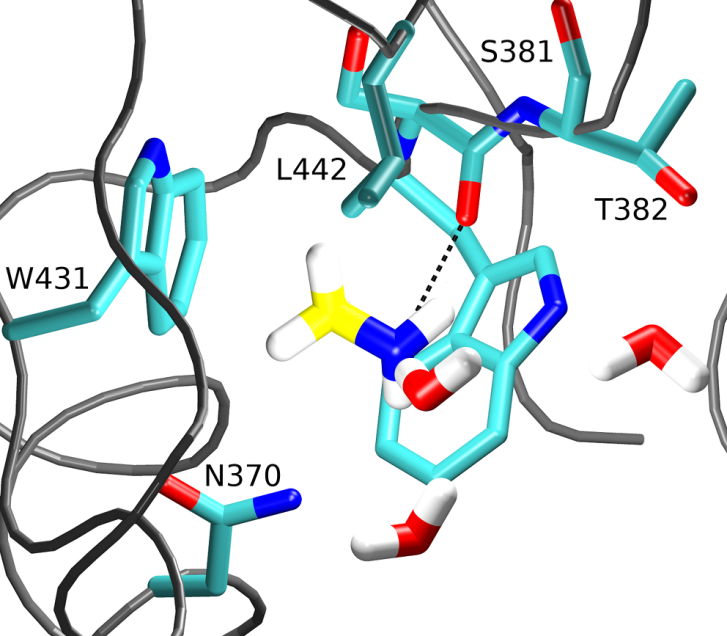

Figure S3. In simulations of the free YTH domain, methylamine bound stably in the m^6^A_3_/A_3_ binding pocket only when water molecules were totally excluded from the region between the methylamine and the W431 side-chain.

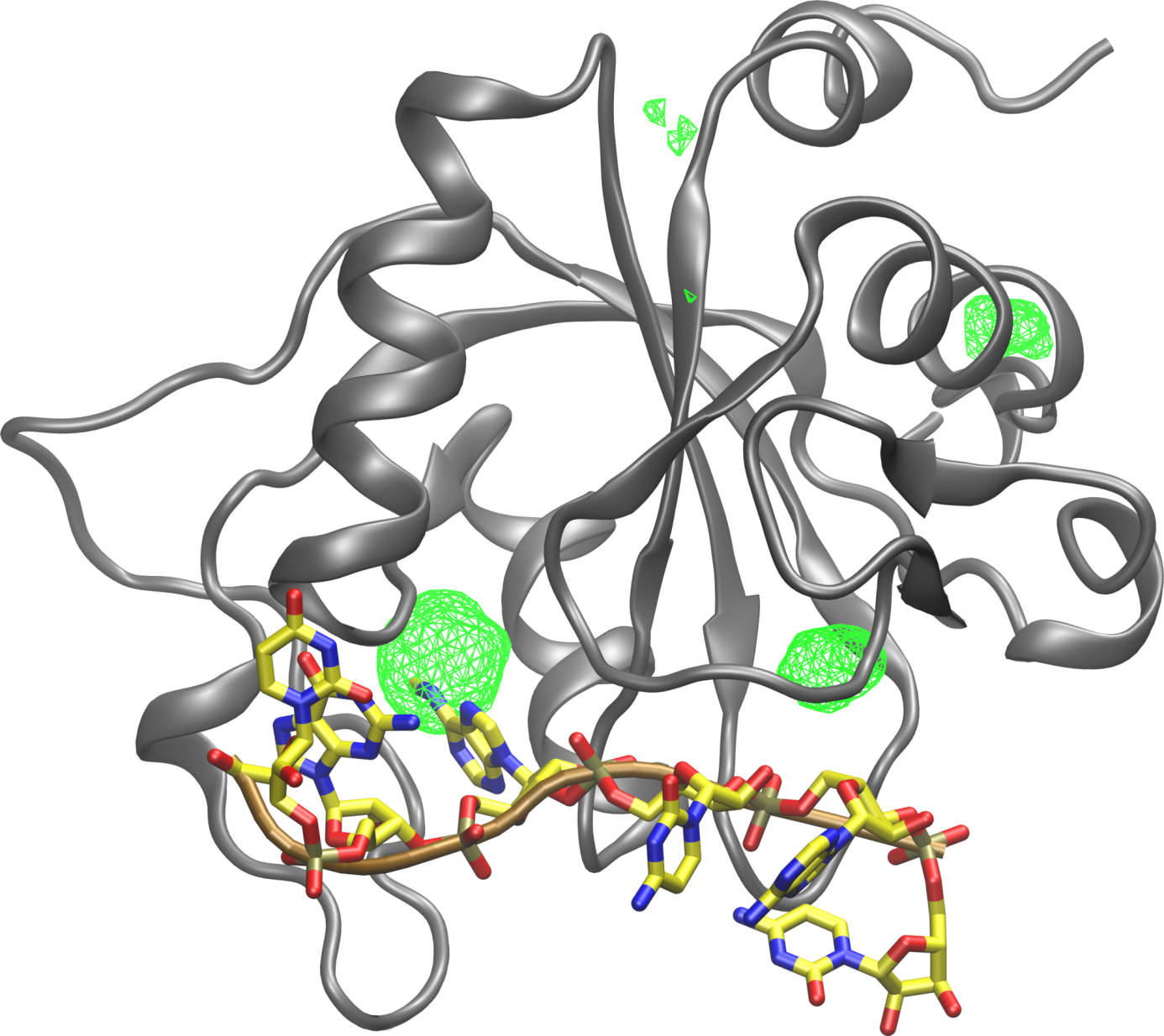

Figure S4. The green wireframe indicates five locations with highest density of methylamines in MD simulations of the free YTH domain. The m^6^A_3_/A_3_ binding pocket had the highest density. The RNA is shown only for comparison and was not included in these simulations.

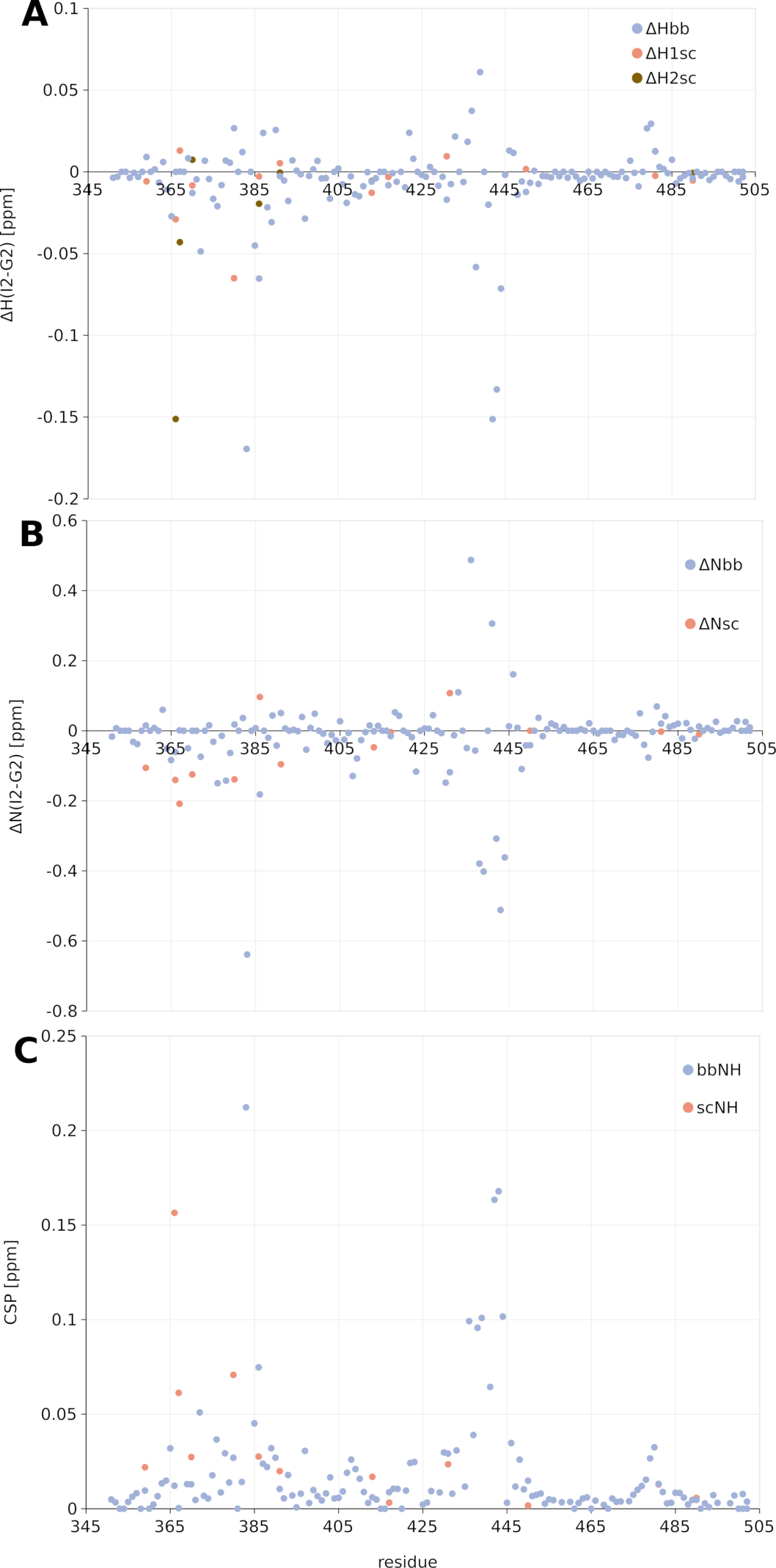

Figure S5. Backbone and side-chain chemical shift perturbations for signals of 1:1 G_2_ RNA versus I_2_ RNA complexed with YTH, plotted against the protein residue number. (**A**) ^1^H chemical shift differences. Side chain ^1^H signals correspond to Asn (Hδ21 and Hδ22), Gln (Hε21 and Hε22), Arg (Hε), and Trp (Hε1). In case of the side chain amines of Asn and Gln, the two separate ^1^H shift changes are shown in orange and brown. (**B**) ^15^N chemical shift differences. Shift changes of the side chain signals of Asn (Nδ2), Gln (Nε2), Arg (Nε), and Trp (Nε1) are shown in orange. (**C**) Combined chemical shift changes calculated using the formula CSP = [(0.2∙ΔN)^2^ + ∑ΔH_i_^2^]^1/2^ where i ranges over all the hydrogen atoms attached to the nitrogen atoms. For side-chain amides of Asn and Gln, this corresponds to two hydrogen atoms.

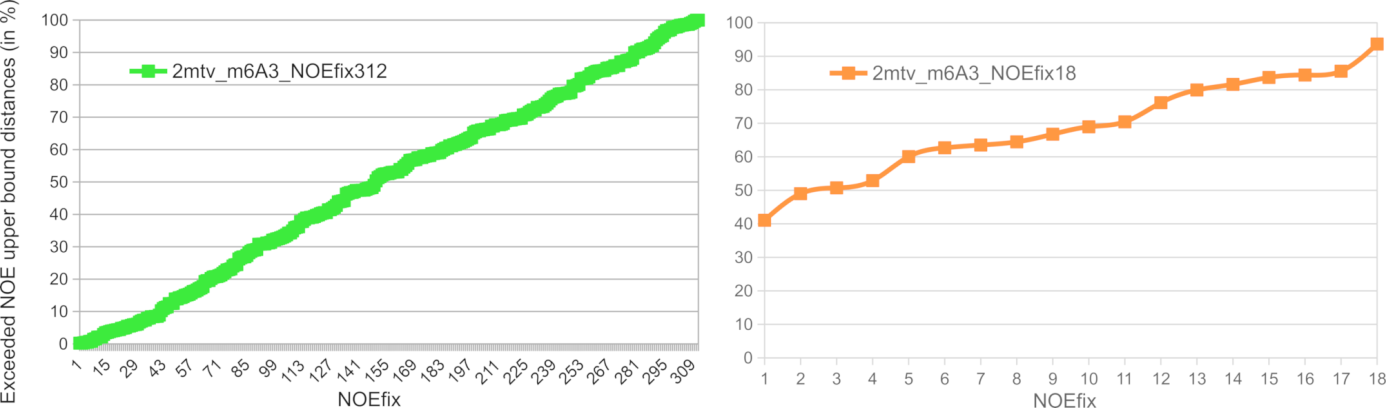

Figure S6. Graphs showing the percentage of simulation frames in which the individual NOEs for which we applied NOEfix exceeded the NOE upper bound distance. The data points were sorted so that from left to right, the curve goes from the least to most often exceeded NOE upper bound. In case of 2mtv_m^6^A_3__NOEfix312 simulation ensemble (left; green), a significant number of the NOE upper bounds were almost never exceeded in simulations, reflecting an undesirable conformational restriction of the system when using large number of NOEfixes. The problem disappeared when only limited number of NOEfixes was used (right) as all targeted distances could exceed the NOE upper bound.

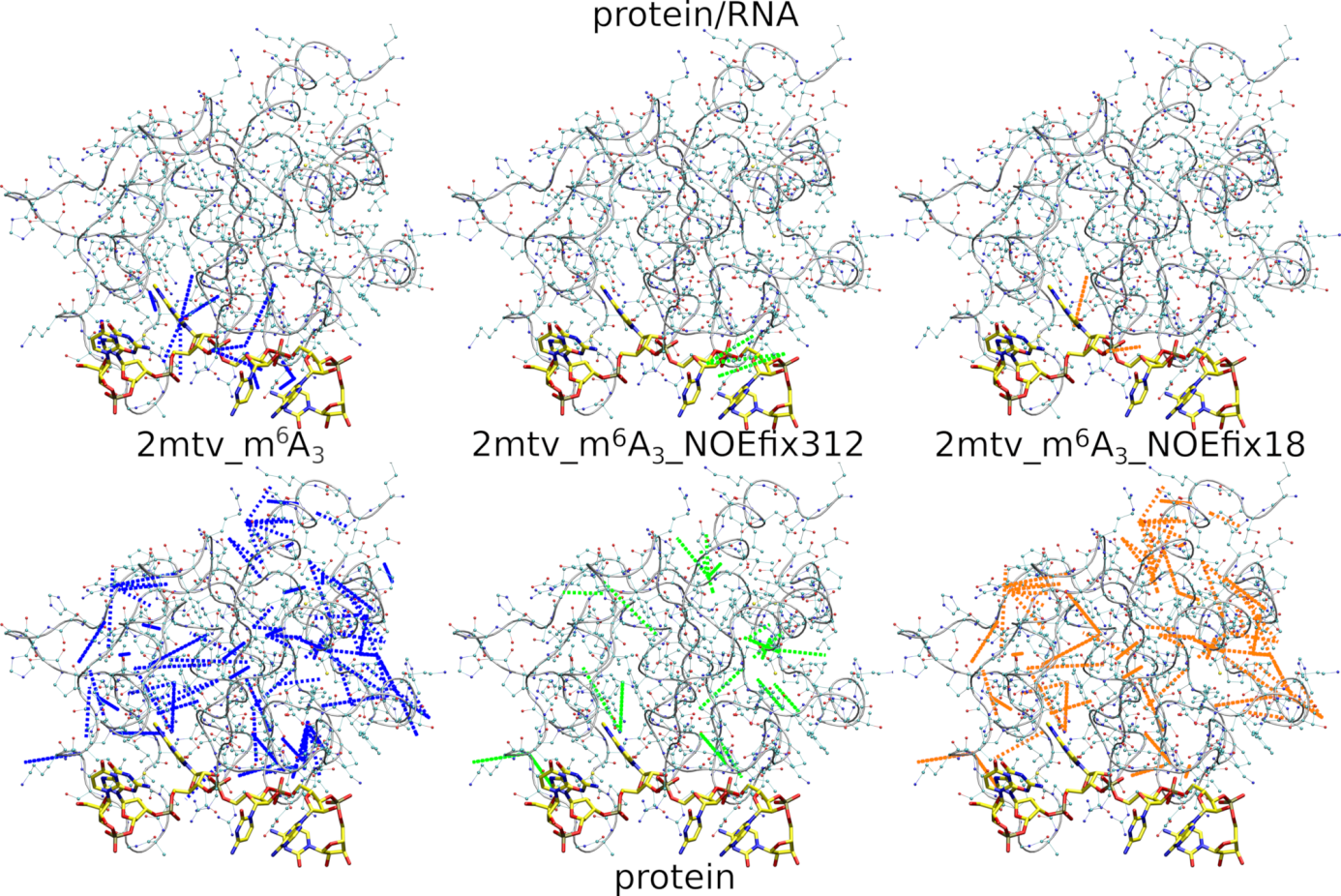
Figure S7. Visualization of the NOE distances violated in the 2mtv_m^6^A_3_ (left; blue lines), 2mtv_m^6^A_3__NOEfix312 (middle; green lines), and 2mtv_m^6^A_3__NOEfix18 (right; orange lines) simulation ensembles. Protein/RNA violations over 0.3 Å and protein/protein violations over 1 Å are visualized on top and bottom, respectively.

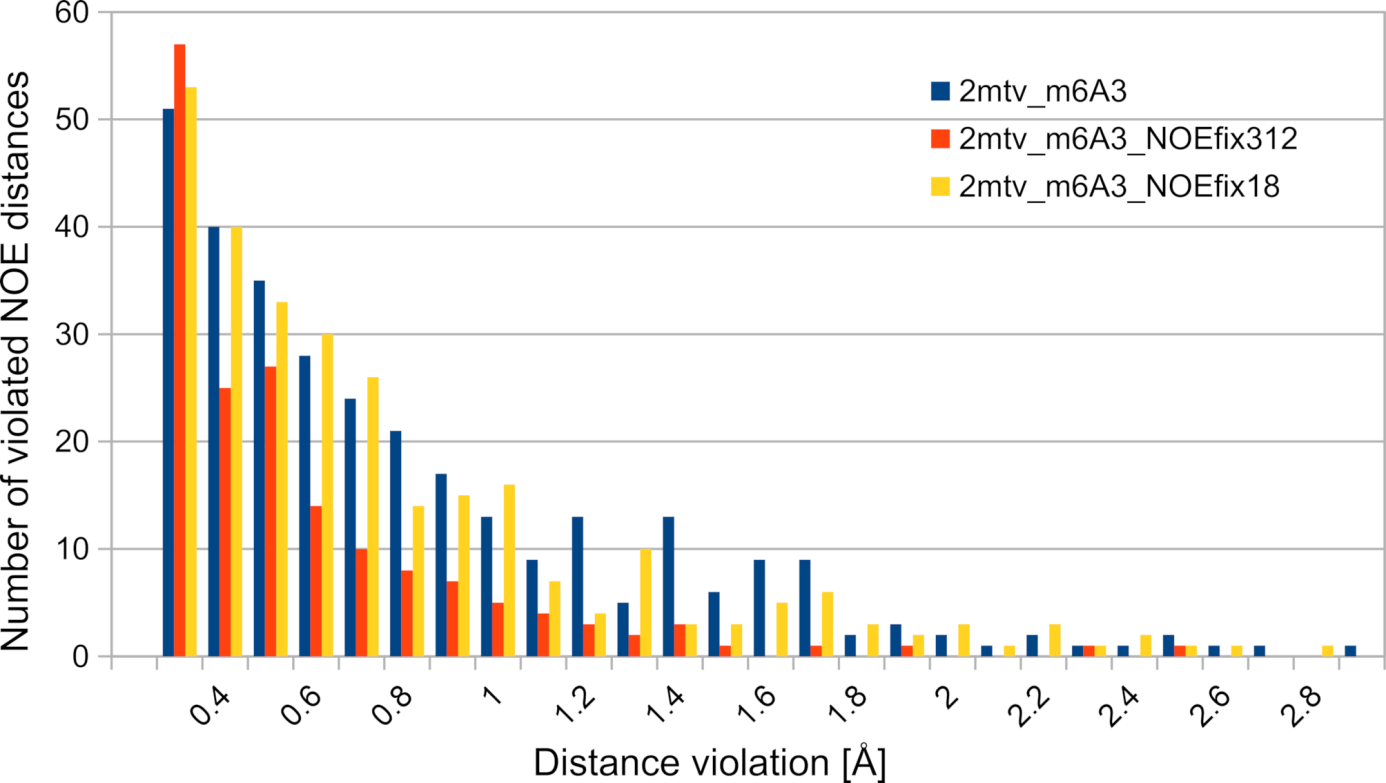

Figure S8. Histograms of the violated NOE distances in standard (where two structure-specific HBfix potentials related to binding of A_5_ and C_6_ nucleotides were utilized; see above) and NOEfix ensembles of the YTH protein/RNA complex simulations. It is evident that NOEfix reduces the overall number of violations, especially violations larger than 1 Å, mainly for the protein-RNA interface (Figure S7).

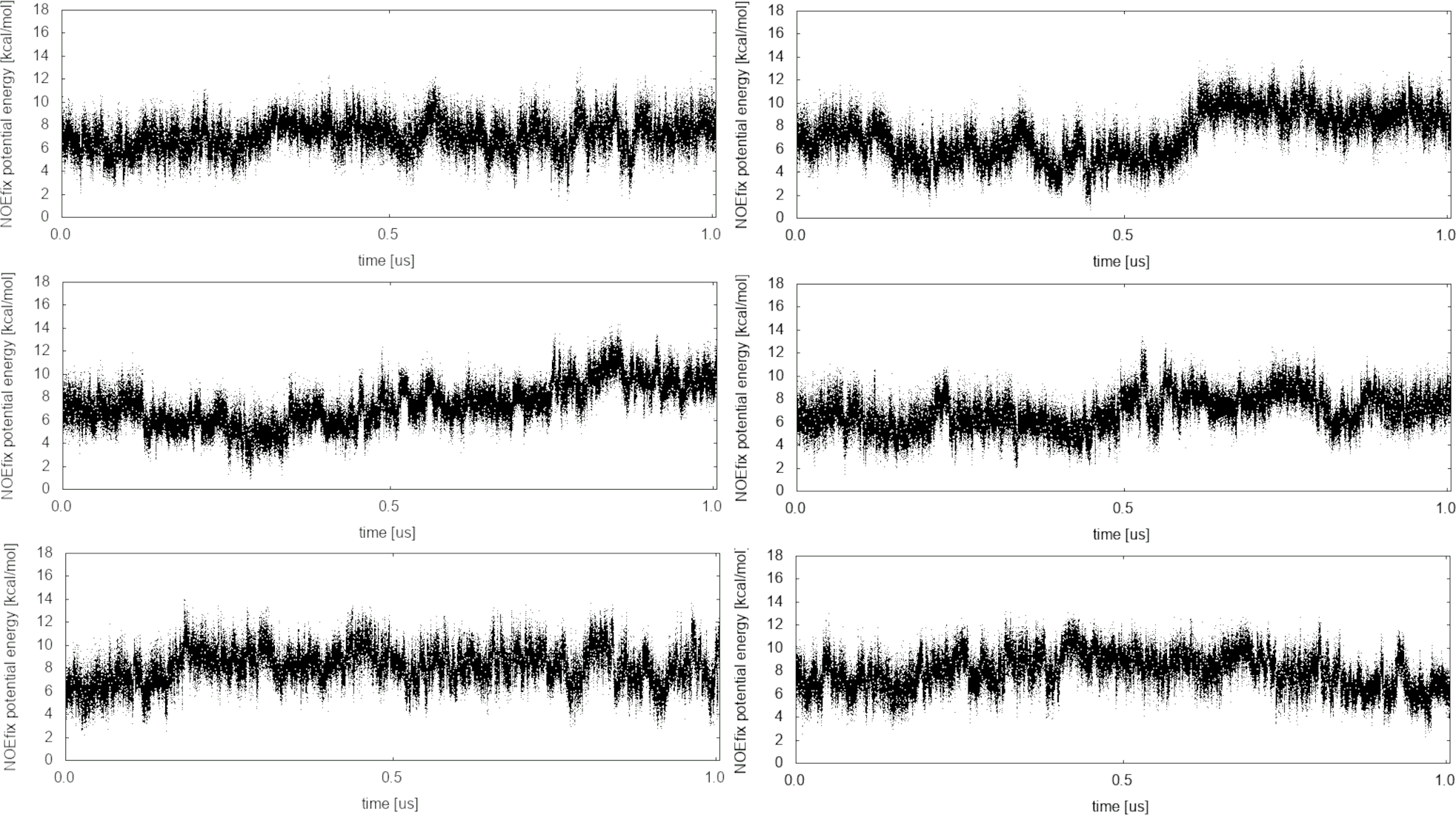

Figure S9. Time development of the cumulative potential energy produced by NOEfix in the individual 2mtv_m^6^A_3__NOEfix18 simulations. 18 NOEfix potentials were used, each with maximum penalty value of 1 kcal/mol. Note that the overall NOEfix potential energy shows only moderate fluctuations which indicates that the simulations are not severely restricted, which is consistent also with other analyses, indicating no spurious suppression of the dynamics.

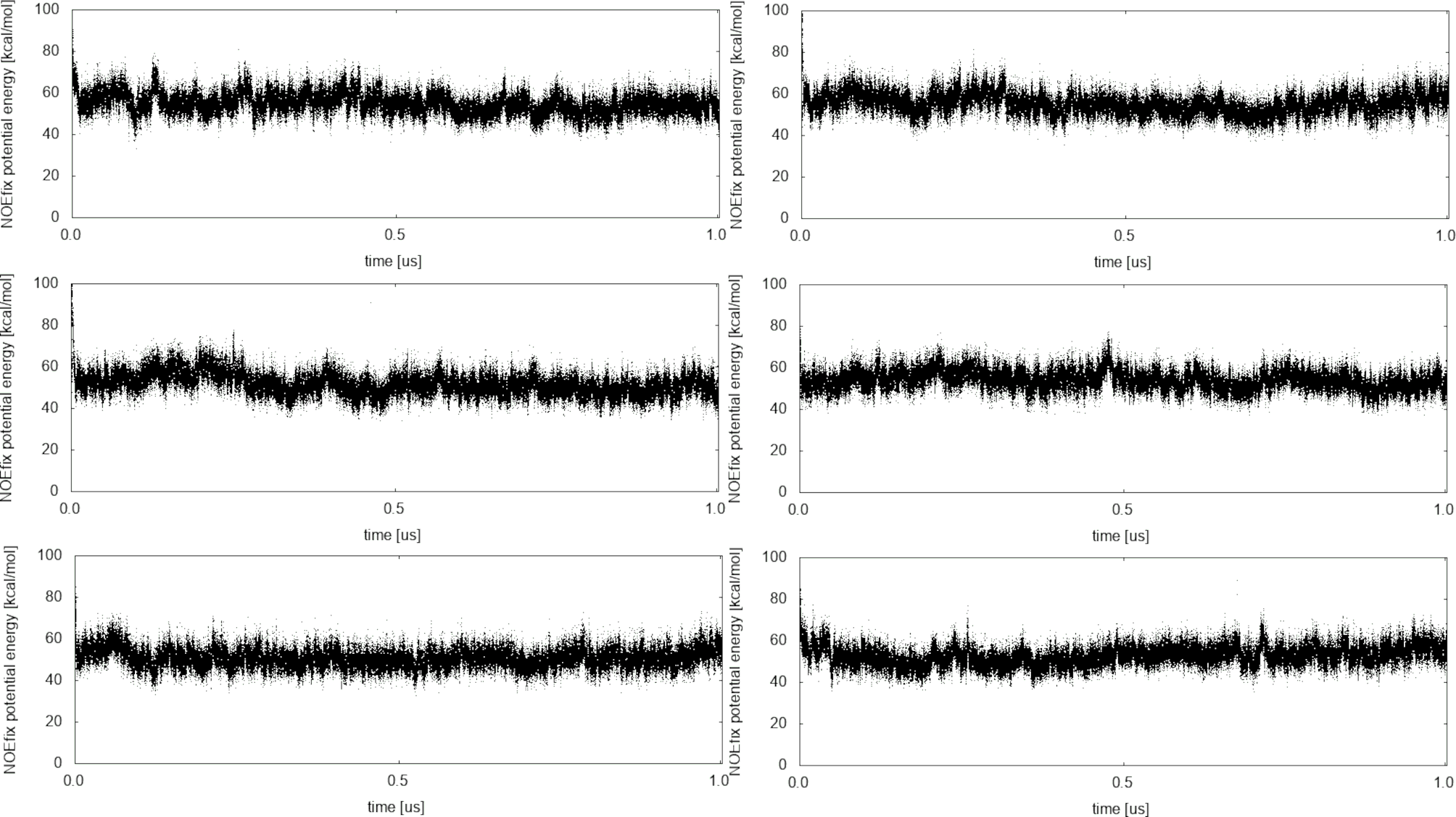

Figure S10. Time development of the cumulative potential energy produced by NOEfix in the individual 2mtv_m^6^A_3__NOEfix312 simulations. 312 NOEfix potentials were used, each with maximum penalty value of 1 kcal/mol.

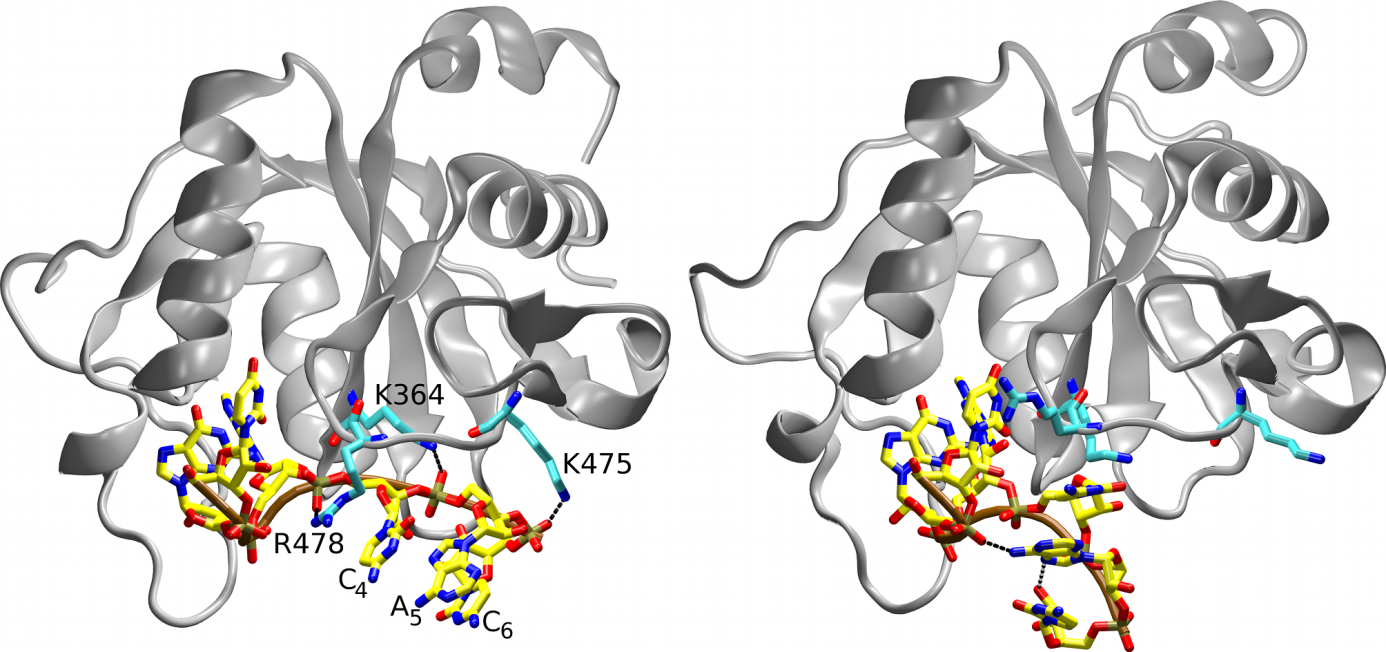

Figure S11. In the preliminary set of simulations, the native protein/RNA H-bond interactions of the C_4_, A_5_, and C_6_ nucleotides (left) were sometimes lost with subsequent formation of (permanent on the simulation time scale) spurious intramolecular RNA interactions (right) that prevented reformation of the protein/RNA interface. The spurious behavior was initiated by fluctuations of H-bonds between the cationic side-chains and the phosphate groups followed by the irreversible RNA compaction. We assume that the RNA self-interaction is the primary force-field issue, as some fluctuations of the protein-RNA H-bonds can be expected. We found several simple ways how to eliminate this behavior for the present system. The problem was entirely fixed by the structure-specific HBfix potential function as well as by the NOEfix; note that the NOEfix simulations did not utilize any HBfix, so the performance of the NOEfix is quite promising. Note also that the image on the right shows just one specific example of spurious RNA self-interactions; different ones were observed in other simulations.

### List of NOE distances (as deposited in the PDB database; 6MA is m^6^A_3_) for which NOEfixes were applied in 2mtv_m^6^A_3__NOEfix18 simulations:

\# 1 U5 H3' 2 G H8 6.00

\# 5 A H8 6 C H1' 5.00

\# 22 LEU QD1 4 C H4' 6

\# 30 ASN HA 3 6MA H2 6

\# 43 LEU HG 3 6MA QH1 6

\# 67 ARG QG 4 C H6 6

\# 68 GLU QB 6 C H5 6

\# 94 PRO HB2 3 6MA H2 5

\# 138 ARG HD2 4 C H3' 6

\# 139 ASP HA 3 6MA H8 6

\# 101 MET QB 2 G H8 4

\# 23 ILE QG1 42 THR QG2 4.3

\# 23 ILE QG1 46 ASN HB3 5.46

\# 24 LYS QD 65 SER H 4.14

\# 27 ASN H 94 PRO HB3 5.5

\# 66 VAL H 69 SER QB 4.84

\# 91 TRP HD1 103 GLY H 5.03

\# 92 VAL H 102 LEU QQD 5.15

### List of NOE distances (as deposited in the PDB database; 6MA is m^6^A_3_) for which NOEfixes were applied in 2mtv_m^6^A_3__NOEfix312 simulations:

\# 1 U5 H3' 2 G H8 6.00

\# 5 A H8 6 C H1' 5.00

\# 22 LEU QD1 4 C H4' 6

\# 30 ASN HA 3 6MA H2 6

\# 43 LEU HG 3 6MA QH1 6

\# 67 ARG QG 4 C H6 6

\# 68 GLU QB 6 C H5 6

\# 68 GLU QG 6 C H5 6

\# 94 PRO HB2 3 6MA H2 5

\# 94 PRO QG 3 6MA H2 3

\# 101 MET QB 2 G H8 4

\# 138 ARG HD2 4 C H3' 6

\# 138 ARG HD2 4 C H4' 4

\# 139 ASP HA 3 6MA H8 6

\# 8 THR HB 12 LYS H 4.14

\# 11 LEU HG 111 ILE QG2 3.73

\# 11 LEU QD2 15 LEU H 4.49

\# 13 SER H 16 GLN HE21 5.29

\# 14 VAL H 61 ILE QD1 5.18

\# 14 VAL HA 18 ALA QB 2.96

\# 14 VAL HA 77 ARG QD 4.9

\# 14 VAL QQG 77 ARG HB2 3.53

\# 14 VAL QQG 77 ARG QG 3.65

\# 15 LEU H 18 ALA QB 4.4

\# 15 LEU QD1 152 CYS H 4.31

\# 15 LEU QQD 150 GLN H 5.44

\# 15 LEU QQD 150 GLN HA 4.92

\# 15 LEU QQD 152 CYS H 3.69

\# 16 GLN H 58 ARG QD 5.28

\# 16 GLN HB2 58 ARG QD 4.83

\# 16 GLN HE22 58 ARG QD 5.04

\# 16 GLN HG2 18 ALA H 5.06

\# 16 GLN HG2 58 ARG QD 4.76

\# 16 GLN HG3 18 ALA H 5.06

\# 16 GLN HG3 58 ARG QD 4.76

\# 16 GLN QG 18 ALA H 4.33

\# 16 GLN QG 58 ARG QD 4.12

\# 17 ASP H 145 LEU QQD 3.44

\# 17 ASP H 58 ARG QD 4.82

\# 17 ASP QB 145 LEU QQD 4.16

\# 18 ALA H 145 LEU HG 5.3

\# 18 ALA H 145 LEU QQD 2.86

\# 18 ALA H 57 ALA QB 5.5

\# 18 ALA H 58 ARG HB3 4.46

\# 18 ALA H 58 ARG QD 5.09

\# 18 ALA QB 57 ALA QB 4.44

\# 18 ALA QB 58 ARG HB3 3.95

\# 18 ALA QB 59 SER HB2 3.17

\# 19 ARG H 145 LEU QQD 4.83

\# 20 PHE H 143 ILE QG2 4.64

\# 20 PHE H 20 PHE QE 4.54

\# 20 PHE H 61 ILE HB 4.97

\# 20 PHE HB2 63 ILE QD1 3.71

\# 20 PHE QE 150 GLN H 5.5

\# 21 PHE H 50 LEU QQD 5.01

\# 21 PHE HA 60 VAL QQG 5.21

\# 22 LEU QD1 138 ARG H 5.5

\# 23 ILE HA 46 ASN HD21 5.5

\# 23 ILE HB 40 TRP HZ2 5

\# 23 ILE QD1 46 ASN HD21 4.58

\# 23 ILE QG1 42 THR QG2 4.3

\# 23 ILE QG1 46 ASN HB3 5.46

\# 23 ILE QG2 42 THR HA 5.5

\# 23 ILE QG2 42 THR QG2 4.45

\# 24 LYS HG2 140 GLY H 5.21

\# 24 LYS QD 65 SER H 4.14

\# 26 ASN H 94 PRO HB3 5.5

\# 26 ASN H 94 PRO HG3 4.36

\# 26 ASN H 97 MET QE 4.77

\# 26 ASN HB2 94 PRO HB2 4.52

\# 26 ASN HB3 94 PRO HB2 4.52

\# 26 ASN QB 94 PRO HB2 3.95

\# 27 ASN H 94 PRO HB3 5.5

\# 27 ASN HA 28 HIST HD2 5.03

\# 28 HIST HB2 29 GLU QB 5.25

\# 28 HIST HB3 31 VAL QG1 4.97

\# 28 HIST HD2 66 VAL QQG 3.95

\# 28 HIST HE1 66 VAL HB 5.5

\# 28 HIST HE1 66 VAL QG1 4.11

\# 28 HIST HE1 66 VAL QG2 4.11

\# 28 HIST HE1 69 SER H 5.04

\# 30 ASN HD21 33 LEU QQD 4.37

\# 30 ASN HD21 89 ILE QG2 6

\# 31 VAL H 64 PHE HB3 6

\# 31 VAL QG2 74 GLY H 5.5

\# 31 VAL QQG 110 TRP HZ3 4.07

\# 31 VAL QQG 64 PHE H 5.44

\# 31 VAL QQG 66 VAL H 4.58

\# 31 VAL QQG 73 GLN HA 4.24

\# 31 VAL QQG 74 GLY H 5

\# 33 LEU H 35 LYS QG 4.93

\# 33 LEU QD1 90 HIST HB2 4.65

\# 33 LEU QQD 89 ILE HA 4.27

\# 33 LEU QQD 90 HIST HB2 3.82

\# 33 LEU QQD 90 HIST HB3 4.22

\# 34 ALA H 110 TRP HZ2 4.67

\# 34 ALA HA 89 ILE QG2 4.87

\# 34 ALA QB 89 ILE QG2 4.72

\# 35 LYS HA 39 VAL H 4.69

\# 35 LYS HB2 110 TRP HE1 4.97

\# 35 LYS QB 110 TRP HE1 4.27

\# 35 LYS QG 36 ALA QB 4.87

\# 35 LYS QG 37 LYS H 4.96

\# 36 ALA H 37 LYS QG 4.22

\# 39 VAL H 89 ILE QD1 4.74

\# 39 VAL H 89 ILE QG2 5.23

\# 39 VAL H 105 VAL QQG 5.17

\# 39 VAL HB 106 PHE QD 4.72

\# 39 VAL QQG 89 ILE HA 4.98

\# 40 TRP HA 91 TRP HE1 5.5

\# 40 TRP HD1 62 LEU QQD 3.32

\# 42 THR H 50 LEU QQD 5.15

\# 43 LEU H 47 GLU QG 5.34

\# 43 LEU H 101 MET QB 5.27

\# 43 LEU HB2 45 VAL HB 4.97

\# 45 VAL QQG 49 LYS QG 3.35

\# 46 ASN HB3 47 GLU QG 5.34

\# 46 ASN HB3 49 LYS H 5.11

\# 48 LYS H 51 ASN HD22 5.5

\# 48 LYS QE 49 LYS H 4.73

\# 49 LYS QE 140 GLY HA2 3.37

\# 50 LEU H 62 LEU QQD 5.34

\# 50 LEU H 78 LEU QQD 5.2

\# 50 LEU HA 62 LEU QQD 4.17

\# 50 LEU QQD 62 LEU H 5.44

\# 50 LEU QQD 62 LEU HA 4.55

\# 50 LEU QQD 63 ILE H 5.44

\# 51 ASN H 52 LEU HG 4.72

\# 51 ASN H 78 LEU QQD 4.42

\# 53 ALA H 55 ARG QD 5.34

\# 53 ALA H 60 VAL QQG 3.98

\# 53 ALA H 78 LEU QQD 5.29

\# 53 ALA QB 60 VAL HA 4.53

\# 54 PHE H 55 ARG QD 4.5

\# 54 PHE HA 55 ARG HD2 5.5

\# 54 PHE HA 55 ARG HD3 5.5

\# 55 ARG H 57 ALA H 4.12

\# 55 ARG H 60 VAL QQG 4.59

\# 55 ARG HB3 57 ALA H 5.5

\# 55 ARG HD2 56 SER H 5.05

\# 55 ARG HD3 56 SER H 5.05

\# 55 ARG QD 57 ALA H 5.34

\# 61 ILE QD1 109 ASP QB 5.34

\# 61 ILE QD1 75 PHE QE 3.86

\# 62 LEU QQD 80 SER H 5.44

\# 63 ILE QG2 72 PHE H 5.5

\# 63 ILE QG2 72 PHE HB2 4.37

\# 64 PHE HZ 110 TRP HE1 5

\# 66 VAL H 69 SER QB 4.84

\# 66 VAL H 73 GLN HE22 4.61

\# 66 VAL H 73 GLN QE2 3.76

\# 66 VAL HB 69 SER HA 5.5

\# 66 VAL HB 69 SER QB 4

\# 68 GLU HA 70 GLY QA 5.34

\# 69 SER H 73 GLN QE2 5.34

\# 69 SER HA 73 GLN QE2 5.34

\# 69 SER HB2 73 GLN HE21 4.86

\# 69 SER HB2 73 GLN HE22 4.86

\# 69 SER HB3 73 GLN HE21 4.86

\# 69 SER HB3 73 GLN HE22 4.86

\# 69 SER QB 73 GLN QE2 3.58

\# 70 GLY H 73 GLN QE2 5.31

\# 71 LYS H 73 GLN HE21 4.55

\# 71 LYS H 73 GLN HE22 4.55

\# 71 LYS H 73 GLN QE2 3.74

\# 71 LYS HD3 115 GLU H 5.5

\# 71 LYS HD3 116 LEU H 3.93

\# 71 LYS HE3 117 PRO HB2 5.5

\# 71 LYS HG2 117 PRO HD2 6

\# 71 LYS HG2 72 PHE HB3 5.5

\# 71 LYS HG3 117 PRO HB2 4.49

\# 72 PHE QD 134 VAL QQG 4.18

\# 73 GLN H 115 GLU QB 5.14

\# 75 PHE H 155 PHE QD 5.18

\# 75 PHE H 155 PHE QE 4.51

\# 76 ALA QB 111 ILE HA 5.46

\# 76 ALA QB 79 SER H 5.5

\# 78 LEU HA 80 SER H 3.56

\# 78 LEU QQD 106 PHE H 4.6

\# 80 SER H 108 ILE H 5.5

\# 80 SER H 108 ILE QG2 4.23

\# 82 SER H 104 GLY HA2 6

\# 83 HIST H 105 VAL HB 4.08

\# 84 HIST HD2 103 GLY HA3 5.5

\# 84 HIST HD2 105 VAL H 5.3

\# 89 ILE H 105 VAL QQG 3.27

\# 89 ILE HB 91 TRP HB2 5.33

\# 89 ILE HG13 105 VAL HA 5.07

\# 89 ILE QG2 91 TRP HE3 3.62

\# 89 ILE QG2 92 VAL H 5.5

\# 91 TRP HD1 103 GLY H 5.03

\# 91 TRP HE1 105 VAL HA 4.41

\# 92 VAL H 102 LEU QQD 5.15

\# 92 VAL QQG 94 PRO HA 3.92

\# 94 PRO HD3 97 MET QE 4.04

\# 97 MET HA 101 MET QE 3.28

\# 98 SER H 102 LEU HG 5.5

\# 99 ALA QB 100 LYS QD 3.26

\# 102 LEU H 104 GLY H 5.5

\# 102 LEU HA 104 GLY H 4.43

\# 102 LEU HA 104 GLY HA3 5.22

\# 103 GLY H 105 VAL H 5.5

\# 104 GLY H 105 VAL H 3.53

\# 105 VAL QQG 107 LYS QE 4.69

\# 106 PHE H 108 ILE QG2 4.43

\# 109 ASP H 111 ILE QD1 4.86

\# 109 ASP HA 111 ILE QD1 5.23

\# 110 TRP HZ3 113 ARG QD 5.34

\# 111 ILE HG12 112 CYS H 4.09

\# 113 ARG H 158 ASP H 5.5

\# 113 ARG QG 161 ILE QD1 3.88

\# 113 ARG QG 161 ILE QG2 4.27

\# 114 ARG H 156 PRO HG2 5.03

\# 116 LEU HG 154 LEU H 5.5

\# 116 LEU HG 154 LEU QD1 3.22

\# 116 LEU HG 155 PHE HB2 5.5

\# 116 LEU QD2 117 PRO HG2 5.18

\# 116 LEU QD2 120 LYS H 4.35

\# 116 LEU QQD 119 THR QG2 4.45

\# 116 LEU QQD 156 PRO HB2 5.44

\# 117 PRO HG2 119 THR HB 4.79

\# 117 PRO HG3 119 THR QG2 4.55

\# 119 THR QG2 120 LYS HA 4.01

\# 119 THR QG2 120 LYS QD 4.04

\# 119 THR QG2 120 LYS QE 4.17

\# 119 THR QG2 120 LYS QG 2.55

\# 119 THR QG2 121 SER H 4.35

\# 120 LYS HA 122 ALA H 3.93

\# 120 LYS QB 154 LEU QD1 3.6

\# 121 SER H 134 VAL QQG 4.76

\# 121 SER H 151 LEU QQD 4.88

\# 121 SER H 154 LEU QD1 4

\# 121 SER HB3 134 VAL QG2 5

\# 121 SER QB 134 VAL QQG 3.07

\# 122 ALA H 124 LEU H 4.11

\# 122 ALA H 154 LEU QD2 5.27

\# 124 LEU QQD 147 CYS H 5.34

\# 124 LEU QQD 150 GLN H 4.19

\# 125 THR QG2 133 PRO HD3 3.35

\# 125 THR QG2 135 LYS H 6

\# 127 PRO HA 128 TRP HD1 5.17

\# 127 PRO HB2 131 HIST HE1 3.76

\# 127 PRO HB3 128 TRP HD1 5.03

\# 127 PRO HG2 128 TRP HE1 4.11

\# 127 PRO HG3 128 TRP HD1 4.72

\# 128 TRP HA 128 TRP HD1 2.81

\# 128 TRP HE1 144 GLU HG3 4.12

\# 128 TRP HE1 144 GLU QG 3.62

\# 128 TRP HE3 142 GLU HA 4.79

\# 128 TRP HH2 145 LEU H 4.23

\# 128 TRP HH2 145 LEU QB 5.31

\# 128 TRP HZ2 144 GLU HA 4.45

\# 128 TRP HZ2 144 GLU HG2 4.6

\# 128 TRP HZ2 144 GLU HG3 4.6

\# 128 TRP HZ2 144 GLU QG 3.77

\# 128 TRP HZ2 145 LEU H 4.53

\# 128 TRP HZ3 142 GLU HG3 3.8

\# 128 TRP HZ3 142 GLU QG 3.29

\# 130 GLU H 141 GLN QE2 5.34

\# 130 GLU H 141 GLN QG 5.34

\# 130 GLU HA 132 LYS QB 5.34

\# 132 LYS HA 136 ILE QD1 4.01

\# 133 PRO HB3 136 ILE H 4.79

\# 133 PRO HB3 136 ILE QD1 4.46

\# 133 PRO HD3 136 ILE QD1 3.45

\# 134 VAL HA 151 LEU QQD 4.92

\# 135 LYS H 136 ILE HG13 4.47

\# 136 ILE QG2 138 ARG H 5

\# 141 GLN HB3 143 ILE H 5.5

\# 141 GLN HB3 143 ILE HG12 4.71

\# 143 ILE QG2 149 THR H 4.38

\# 144 GLU HB3 146 GLU HG3 4.31

\# 145 LEU QD1 147 CYS H 5.5

\# 146 GLU H 149 THR HB 4.88

\# 146 GLU HA 149 THR QG2 3.4

\# 146 GLU HB2 149 THR HB 4.59

\# 146 GLU HG3 149 THR H 5.5

\# 147 CYS H 150 GLN QG 5.02

\# 147 CYS QB 150 GLN QG 5.1

\# 148 GLY H 149 THR HB 4.69

\# 148 GLY H 151 LEU HG 4.76

\# 148 GLY H 151 LEU QD1 5.5

\# 148 GLY HA2 151 LEU HG 4.05

\# 148 GLY HA3 151 LEU HG 4.29

\# 149 THR H 150 GLN QG 4.29

\# 149 THR QG2 150 GLN H 3.17

\# 149 THR QG2 150 GLN HA 3.46

\# 149 THR QG2 150 GLN HE21 3.95

\# 149 THR QG2 150 GLN HE22 4.84

\# 149 THR QG2 150 GLN HG2 4.55

\# 149 THR QG2 150 GLN HG3 4.55

\# 149 THR QG2 150 GLN QG 3.7

\# 149 THR QG2 151 LEU H 4.77

\# 150 GLN H 153 LEU QD2 5.5

\# 151 LEU H 153 LEU QD2 6

\# 151 LEU H 154 LEU QD2 4.2

\# 151 LEU HA 154 LEU QD2 3.01

\# 151 LEU HB2 155 PHE HZ 5.5

\# 151 LEU HB3 154 LEU QD1 3.19

\# 151 LEU HB3 155 PHE QD 4.9

\# 152 CYS H 154 LEU QD1 5.23

\# 152 CYS H 155 PHE H 5.14

\# 153 LEU H 154 LEU HG 4.03

\# 153 LEU H 154 LEU QD1 4.79

\# 153 LEU H 154 LEU QD2 4.61

\# 154 LEU H 154 LEU QD2 3.41

\# 154 LEU HG 155 PHE H 3.78

\# 154 LEU QD1 155 PHE H 3.99

\# 157 PRO QG 159 GLU H 3.82

\# 157 PRO QG 159 GLU QB 3.22

\# 158 ASP QB 161 ILE HB 4.15

\# 160 SER H 162 ASP H 4.77
